# Mechanistic basis of choline import involved in teichoic acids and lipopolysaccharide modification

**DOI:** 10.1101/2021.09.14.460277

**Authors:** Natalie Bärland, Anne-Stéphanie Rueff, Gonzalo Cebrero, Cedric A.J. Hutter, Markus A. Seeger, Jan-Willem Veening, Camilo Perez

**Affiliations:** Biozentrum, University of Basel, 4056 Basel, Switzerland; Department of Fundamental Microbiology, Faculty of Biology and Medicine, University of Lausanne, 1015 Lausanne, Switzerland; Institute of Medical Microbiology, University of Zurich, Zurich, Switzerland; Linkster Therapeutics AG

## Abstract

Phosphocholine molecules decorating bacterial cell wall teichoic acids and outer-membrane lipopolysaccharide have significant roles in adhesion to host cells, immune evasion, and persistence. Bacteria carrying the operon that performs phosphocholine decoration, synthesize phosphocholine after uptake of the choline precursor by LicB, a conserved transporter among divergent species. *Streptococcus pneumoniae* is a prominent pathogen where phosphocholine decoration plays a fundamental role in virulence. Here we present cryo-electron microscopy and crystal structures of *S. pneumoniae* LicB, revealing distinct conformational states and describing architectural and mechanistic elements essential to choline import. Together with *in vitro* and *in vivo* functional characterization, we found that LicB displays proton-coupled import activity and promiscuous selectivity involved in adaptation to choline deprivation conditions, and describe LicB inhibition by synthetic nanobodies (sybodies) and hemicholinium-3. Our results provide novel insights into the molecular mechanism of a key transporter involved in bacterial pathogenesis and establish a basis for inhibition of the phosphocholine modification pathway across bacterial phyla.

---

Teichoic acids (TA) are fundamental biopolymers that make part of the cell wall of Gram-positive bacteria (*1-3*), whereas lipopolysaccharides (LPS) are exclusively found at the outer membrane of Gram-negatives (*4-6*). Although their compositions differ, both TA and LPS function as endotoxins in bacterial pathogens, participate in immune evasion, prevent recognition and opsonization by antibodies, and play important roles in adhesion and colonization (*7, 8*). Decoration of TA and LPS with phosphocholine moieties is among the most impactful cell wall modifications, conferring virulence advantages to multiple pathogens (*9-11*). Gram-positive *Streptococcus pneumoniae* and Gram-negative *Haemophilus influenzae*, both residing in the human respiratory tract, are well characterized pathogens that expose phosphocholine epitopes on their cell surface (*12, 13*). Interactions of phosphocholine with host proteins, such as the platelet activating factor receptor (*14-16*), allow adhesion to the surface of host cells followed by cell invasion (*17, 18*). In addition, *S. pneumoniae* and commensal streptococci use phosphocholine decorated TA as a platform that anchors a great variety of choline binding proteins, which contribute to adherence, colonization and virulence (*19-26*).

Bacteria generally import choline from the extracellular milieu and use it for osmoregulation (*27, 28*). However, some bacteria import choline exclusively for modification of TA or LPS (*10, 29, 30*). The important function of this modification leads to choline auxotrophy in *S. pneumoniae* (*31*), although this pathogen can increase the available extracellular natural pool of choline by processing host phospholipids via surface exposed or secreted phosphodiesterases (*32, 33*). Thus, choline import catalysed by LicB, is an essential trait of pathogens like *S. pneumoniae* (**Fig. 1A**) (*31, 34-37*). LicB is a 32-kDa protein, member of the drug/metabolite transporters (DMT) superfamily, which consists of 14 transporter families, harbouring more than 300 membrane proteins ubiquitously distributed in eukaryotes, bacteria and archaea (*38*). Members of this superfamily display 4 to 10 transmembrane (TM) helices, and are derived from primitive proteins of 4 TM helices similar to EmrE, a member of the small multidrug resistance (SMR) family (*39*).

**Figure 1.**
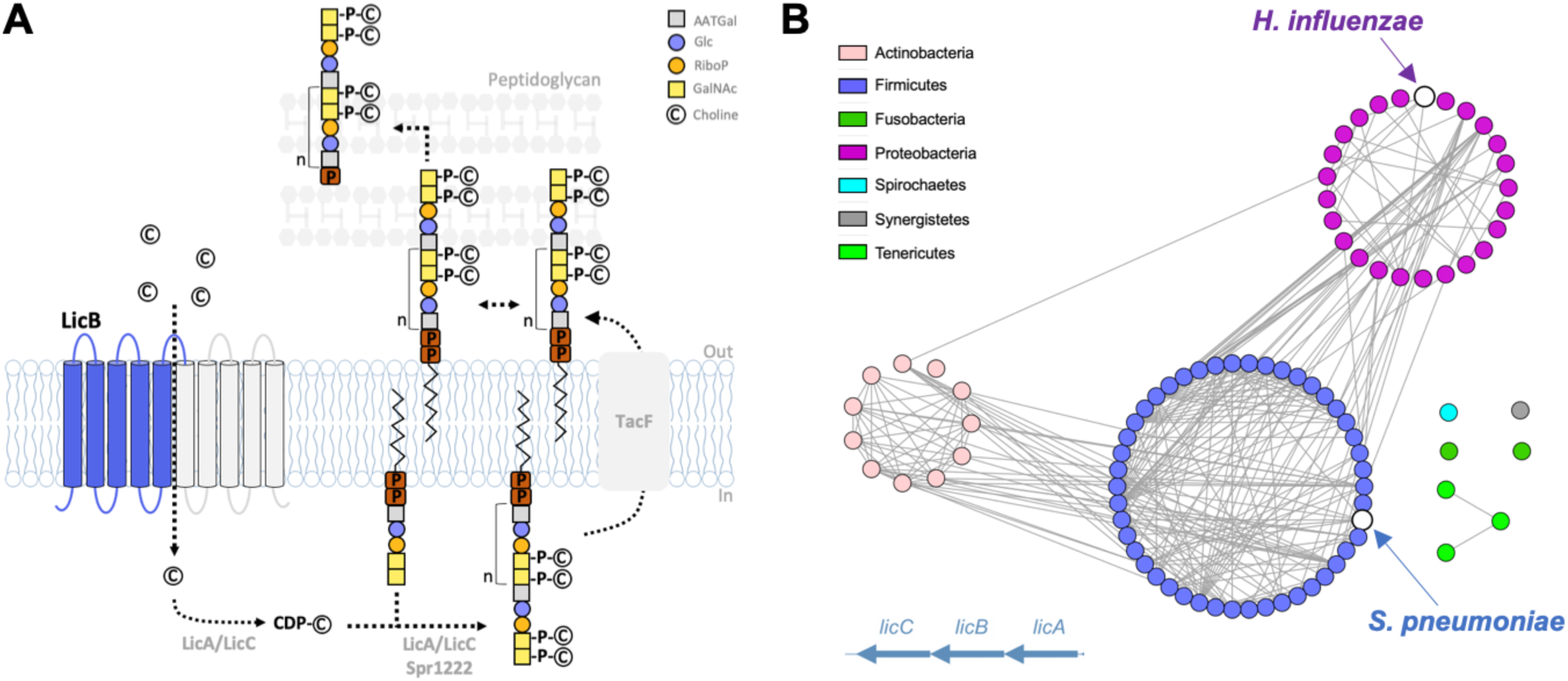
Phosphocholine decoration of teichoic acids and conservation of LicB across divergent bacteria phyla. **A**. Pathway for phosphocholine decoration of teichoic acids (TA) in *S. pneumoniae*. Choline is imported by LicB and activated by LicA and LicC, before being covalently attached as phosphocholine to TA, which are later exposed at the cell wall. Inverted repeats are shown in blue and gray. **B**. Sequence similarity network of bacteria species (nodes) where *lic* operon genes encoding proteins involved in choline uptake (LicB) and activation (LicA and LicC) are conserved. Edges between nodes correspond to an identity of at least 40% among LicB proteins. Sequences are organized according to bacteria phylum (see Fig. S1). *S. pneumoniae* and *H. influenzae* are indicated.

The *licB* gene is located in the *lic* operon which encodes the protein machinery that converts choline into phosphocholine and that decorates TA and LPS (*34, 36, 40*). Inactivation of the *licB* gene leads to non-viable *S. pneumoniae* (*31*), whereas in *H. influenzae* leads to choline uptake deficiency under human nasal airway conditions (*34*). Genes from the *lic* operon can be found in multiple other Gram-positive and Gram-negative bacterial species where phosphocholine decoration of TA, LPS, or other structures occur (*35*). A sequence-similarity network showing *licB* genes sharing high sequence identity (>40%) among different bacterial phyla, demonstrates the important role of this transporter in phosphocholine decoration across Gram-positive and Gram-negative bacteria (**Fig. 1B and Fig. S1**). Despite its clear relevance and potential as drug target, there are no structural or functional studies aiming to characterize the mechanism of LicB or that provide clues on how to inhibit its function.

To elucidate mechanistic elements essential to LicB function, we determined apo-outward-open and choline-bound-occluded structures of *S. pneumoniae* LicB solved by cryo-electron microscopy (cryo-EM) and X-ray crystallography, and performed *in vitro* assays in proteoliposomes and in *S. pneumoniae* cells. We demonstrate that LicB displays promiscuous selectivity involved in adaptation to choline deprivation conditions, and describe architectural elements conserved among divergent bacterial phyla essential for choline binding and proton-coupled import activity. In addition, we describe inhibition of LicB by synthetic nanobodies (sybodies) and hemicholinium-3, thereby establishing the basis for selective inhibition of the phosphocholine decoration pathway.

## Results

### *S. pneumoniae* LicB is a promiscuous high-affinity choline transporter

We characterized the transport activity of LicB reconstituted in proteoliposomes using solid supported membrane (SSM) electrophysiology (*41*) (**Fig. 2A,B**). Transport of choline is electrogenic, as evidenced by a positive current consistent with transport of the positively charged choline into proteoliposomes (**Fig. 2A**). The amplitude of the current measured before reaching electrochemical equilibrium, evidenced by the rapid current decay, is proportional to the initial rate of choline transport. Using this assay, we determined that LicB displays an EC_50_ (apparent K_M_) for choline of 47 ± 15 µM (**Fig. 2B**). In contrast, protein-free liposomes exhibit no transport of choline (**Fig. 2A**). We tested whether LicB can transport alternative compounds with choline-like characteristics. Activation by arsenocholine, a natural compound with a similar composition to choline but containing arsenic instead of nitrogen (*42*), can be transported by LicB as evidenced by positive currents arising from transport into proteoliposomes (**Fig. S2A**). LicB displays lower transport affinity for arsenocholine (EC_50_ = 170 ± 9 µM) (**Fig. 2B**), likely due to the larger radius of the positively charged trimethyl-arsonium group. These results recapitulate prior observations of arsenocholine transport by a choline-transporting variant of the betaine symporter BetP (*43*), and by the choline and betaine transporters OpuB and OpuC, respectively (*44*). In addition, acetylcholine, a positively charged molecule that in contrast to choline carries an ester acetyl instead of a hydroxyl group, is also transported by LicB into proteoliposomes (**Fig. S2B**), albeit with lower affinity (EC_50_ = 740 ± 84 µM) (**Fig. 2B**). Taking together, these results show that LicB is a high-affinity choline transporter that displays promiscuous activity towards other choline-like molecules.

**Figure 2.**
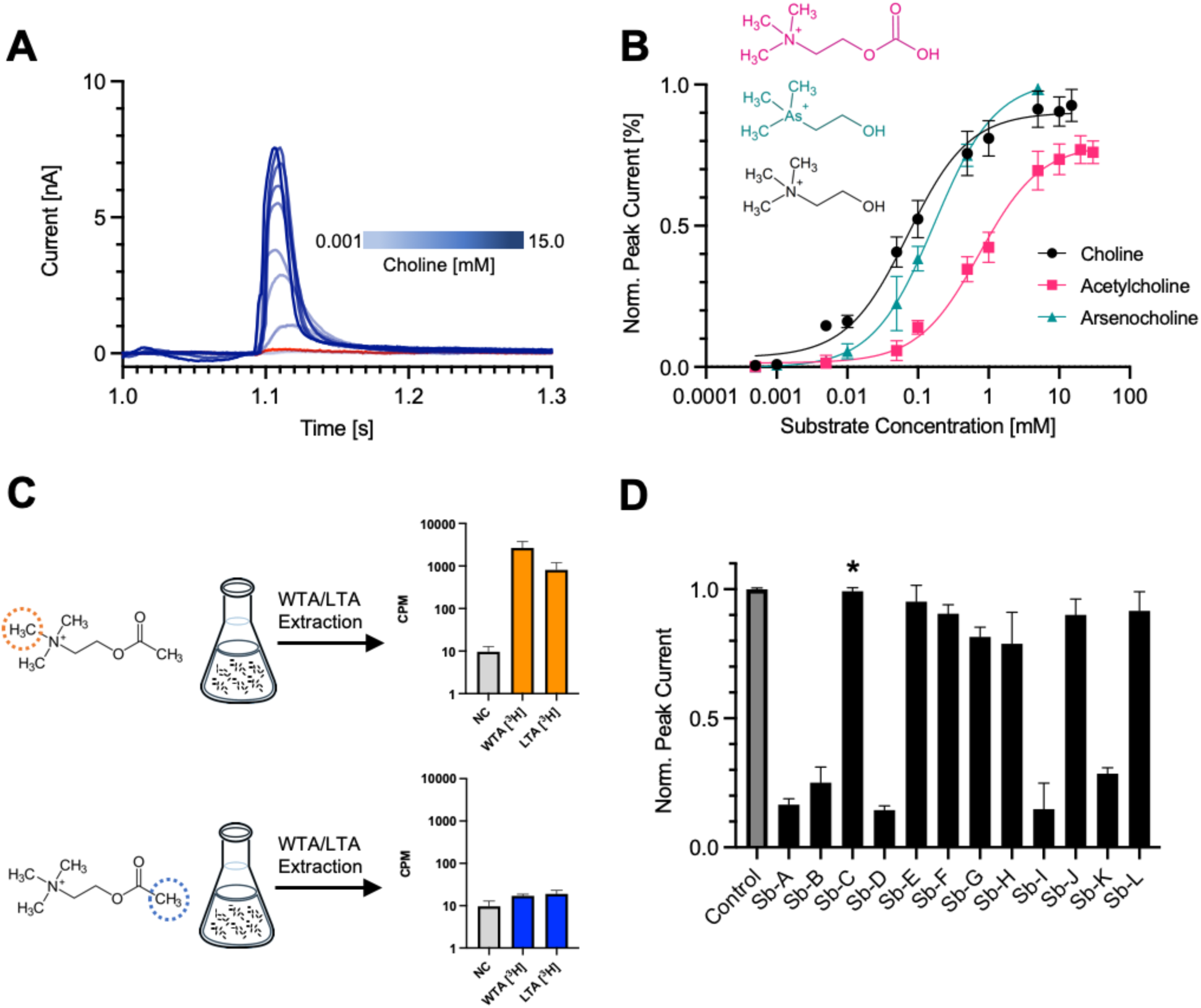
LicB displays promiscuous activity towards choline-like molecules. **A**. SSM-electrophysiology recordings of choline transport in LicB proteoliposomes. Currents recorded between 1-1.3 seconds after addition of choline are shown in blue. Recordings with protein-free liposomes in presence of 5mM choline is shown in red. **B**. Determination of EC_50_ values for choline, arsenocholine and acetylcholine based on SSM recording as shown in A and Fig. S2. **C**. Wall teichoic acids (WTA) and lipoteichoic acids (LTA) extraction from *S. pneumoniae* cells grown in presence of [^3^H]-acetylcholine radioactively labeled at different positions indicated by dotted circles. Histograms show the [^3^H] radioactivity from extracted WTA and LTA. NC, negative control performed with non-radioactive acetylcholine. **D**. Normalized amplitudes of SSM-currents measured in presence of sybodies A to L (500 nM), under 5 mM choline transport conditions based on data shown in Fig. S3. The asterisk denotes Sybody-C, which was used for structural studies. n=3-4 biological replicates, n=2-3 technical replicates, for all experiments.

### Acetylcholine can be processed by *S. pneumoniae* for decoration of teichoic acids

Acetylcholine is a neurotransmitter with function in the central and peripheral nervous system, where it is produced in cholinergic neurons and activates acetylcholine receptors before being processed by acetylcholinesterase. *S. pneumoniae* is one of the major meningitis-causing pathogens in part due to its ability to cross the blood-brain barrier (*45*). During invasion of the central nervous system, *S. pneumoniae* can be exposed to acetylcholine. Thus, in light of the promiscuous transport activity of LicB towards this molecule, we tested whether *S. pneumoniae* can make use of acetylcholine to supply the pathway that carry out decoration of TA. To do this, we grew *S. pneumoniae* D39V cells in choline-reduced media containing acetylcholine isotopically labeled with [^3^H]-methyl at its trimethyl-ammonium group (**Fig. 2C**). After harvesting the cells, we performed extraction of TA and quantification of their radiolabeled content (**Fig. 2C**). Our results show prominent [^3^H] radioactivity in TA extracts, indicating that indeed acetylcholine was used for functionalization of TA.

To demonstrate that acetylcholine has been hydrolyzed by *S. pneumoniae*, and that only its choline moiety was used for decoration of TA, we performed the same experiment but in presence of acetylcholine labeled with [^3^H]-methyl at its acetyl group (**Fig. 2C**). In this case, we expected to see background levels of radioactivity as the choline moiety will remain unlabeled after acetylcholine hydrolysis. Our results show that extracted TA display 100-fold lower levels of [^3^H] radioactivity in comparison to what was observed for extracted TA, when acetylcholine labeled at its trimethyl-ammonium group was used (**Fig. 2C**). These results reveal that *S. pneumoniae* is able to uptake acetylcholine, hydrolyze it, and supply the phosphocholine synthesis pathway for functionalization of TA. Thus, the promiscuous activity of LicB represents an advantage that allows retrieving choline from alternative sources.

### Sybodies inhibit LicB activity but do not affect *S. pneumoniae* growth *in vitro*

We reasoned that due to the small size of LicB (32-kDa), complex formation with a sybody (∼16 kDa) would facilitate structure elucidation by single particle cryo-EM. Thus, we raised sybodies against biotinylated LicB in the presence of 5 mM choline through a combination of ribosome- and phage-display (*46, 47*). Subsequent ELISA screening resulted in the identification of multiple sybodies exhibiting good expression levels and decent behavior on SEC. To characterize the effect of sybodies on choline transport, we reconstituted LicB in proteoliposomes and performed SSM-electrophysiology as described before (*48*). We measured electrogenic transport after activation with 5 mM choline and evaluated peak current amplitudes in the presence and absence of 12 sybodies (**Fig. 2D and Fig. S3**). Five sybodies displayed strong inhibition of LicB transport activity, as evidenced by a decrease of about 80% of the peak currents, whereas seven sybodies did not affect choline electrogenic transport (**Fig. 2D**). Whereas the sequences of the complementarity determining regions (CDRs) of inhibitory sybodies shared high similarity, the CDR regions of non-inhibitory sybodies were more diverse (**Fig. S4**). We further selected a set of three inhibitory sybodies (Sybodies A, B, and D) and one non-inhibitory (Sybody-C), based on their high yields of expression and good behavior during and after purification (**Fig. S5**), and determined their binding kinetics using Grating-coupled interferometry (GCI). This showed that all sybodies bind to LicB with high-affinity displaying binding constants (K_D_) in the range of 30 to 40 nM. Interestingly, binding constants did not differ in the presence of 5 mM choline, indicating that the inhibitory sybodies can bind equally well to choline-bound and substrate-free LicB and are thus not conformationally selective (*49*) (**Fig. S6**).

Due to their small size, strong antigen affinity, low immunogenicity, and easy production, sybodies are strong candidates for development of therapeutics (*50-52*). Thus, we wanted to test whether the inhibitory sybodies A, B and D, were able to affect growth of *S. pneumoniae* due to inhibition of LicB activity. To test this, we grew unencapsulated *S. pneumoniae* D39V cells (strain VL567) in liquid media in presence of the selected inhibitory sybodies, and the non-inhibitory sybody-C as a control, at concentrations ranging from 0.25 µM to 25 µM (**Fig. S7**). Our results indicate that under these experimental conditions, the three inhibitory sybodies tested did not affect *S. pneumoniae* growth. We hypothesize that sybodies might be unable to penetrate the cell wall, or that the remaining LicB choline transport activity, 14% to 28% based on SSM-electrophysiology measurements (**Fig. 2D**), might suffice to supply the pathway for phosphocholine synthesis.

### Structure of outward-open LicB in complex with sybody

We incorporated purified *S. pneumoniae* LicB in nanodiscs containing a mixture of POPG and *E. coli* polar lipid extract (3:1, w:w), and incubated with the non-inhibitory sybody-C during reconstitution (**Fig. S8A**). In contrast to the inhibitory sybodies, sybody-C led to cryo-EM data of better quality. The LicB:Sybody complex structure was solved to a nominal resolution of 3.75 Å (**Fig. 3A, Fig. S8B-D and S9A, and Table 1**). Sybody-C was found bound via its CDR3 region to cytoplasmic loop 4 located between TM4 and TM5 of LicB (**Fig. 3A,B**). The structure revealed a LicB dimer in nanodiscs, with both protomers having the same topological orientation (**Fig. 3A,C**). The dimer interphase involves hydrophobic interactions between aromatic and aliphatic residues located in TM1, TM8, and TM9, and a putative partially resolved POPG lipid molecule, the most prominent phospholipid in the *S. pneumoniae* membrane (*53*) (**Fig. 3C,D and Fig. S9B,C**). The polar headgroup of the lipid faces the extracellular side of the membrane and the aliphatic tails are surrounded mostly by aromatic residues (**Fig. S9B,C**).

**Figure 3.**
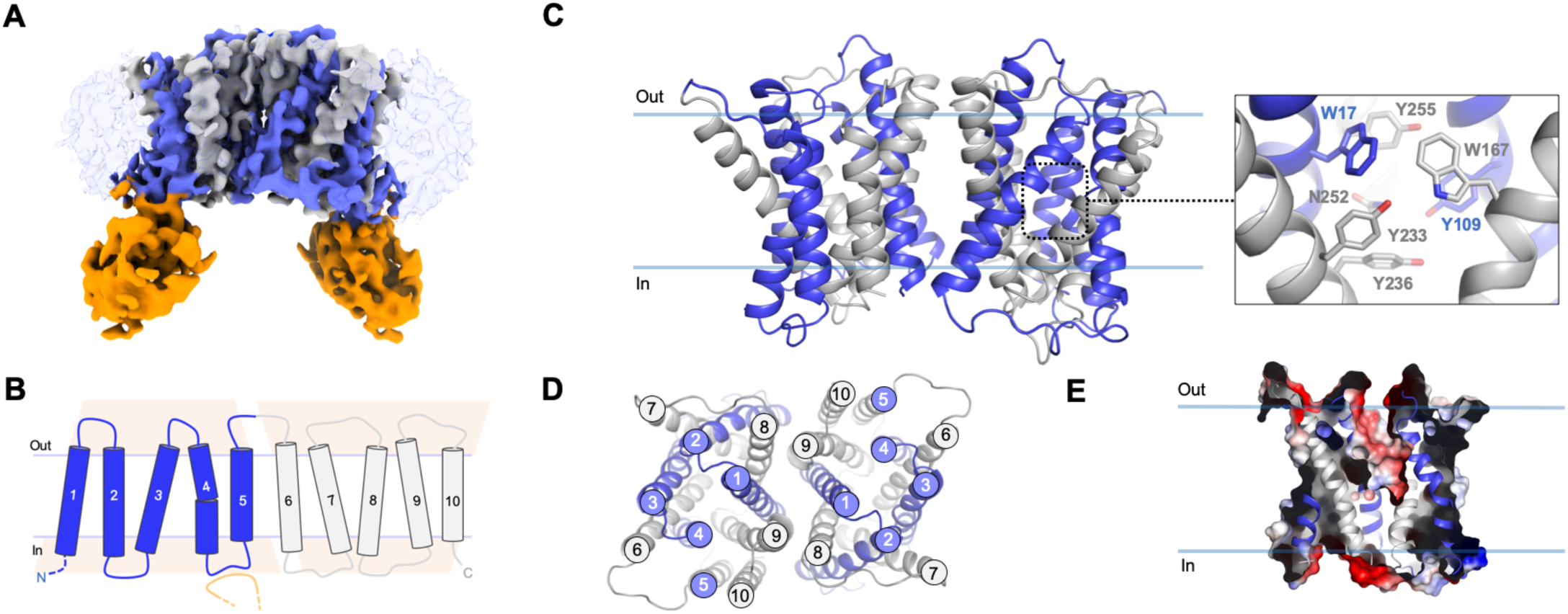
Cryo-EM structure of outward-open *S. pneumoniae* LicB in lipid nanodiscs. **A**. Cryo-EM density of LicB homodimer in nanodiscs with bound sybody at 3.75 Å. Inverted repeats, TM1-5 and TM6-10, in each protomer are colored in blue and grey, respectively. Sybody is colored in orange. The nanodisc is shown as transparent light blue. **B**. Topology of LicB highlighting the inverted antiparallel repeats shown as pink trapezoids. The CDR3 region of the sybody interacting with the loop connecting TM4 and TM5 is shown in orange. **C**. LicB homodimer and central cavity residues. **D**. Top view of LicB homodimer. **E**. Surface electrostatic potential representation of LicB showing the central cavity opening to the extracellular side of the membrane.

**Table 1:**
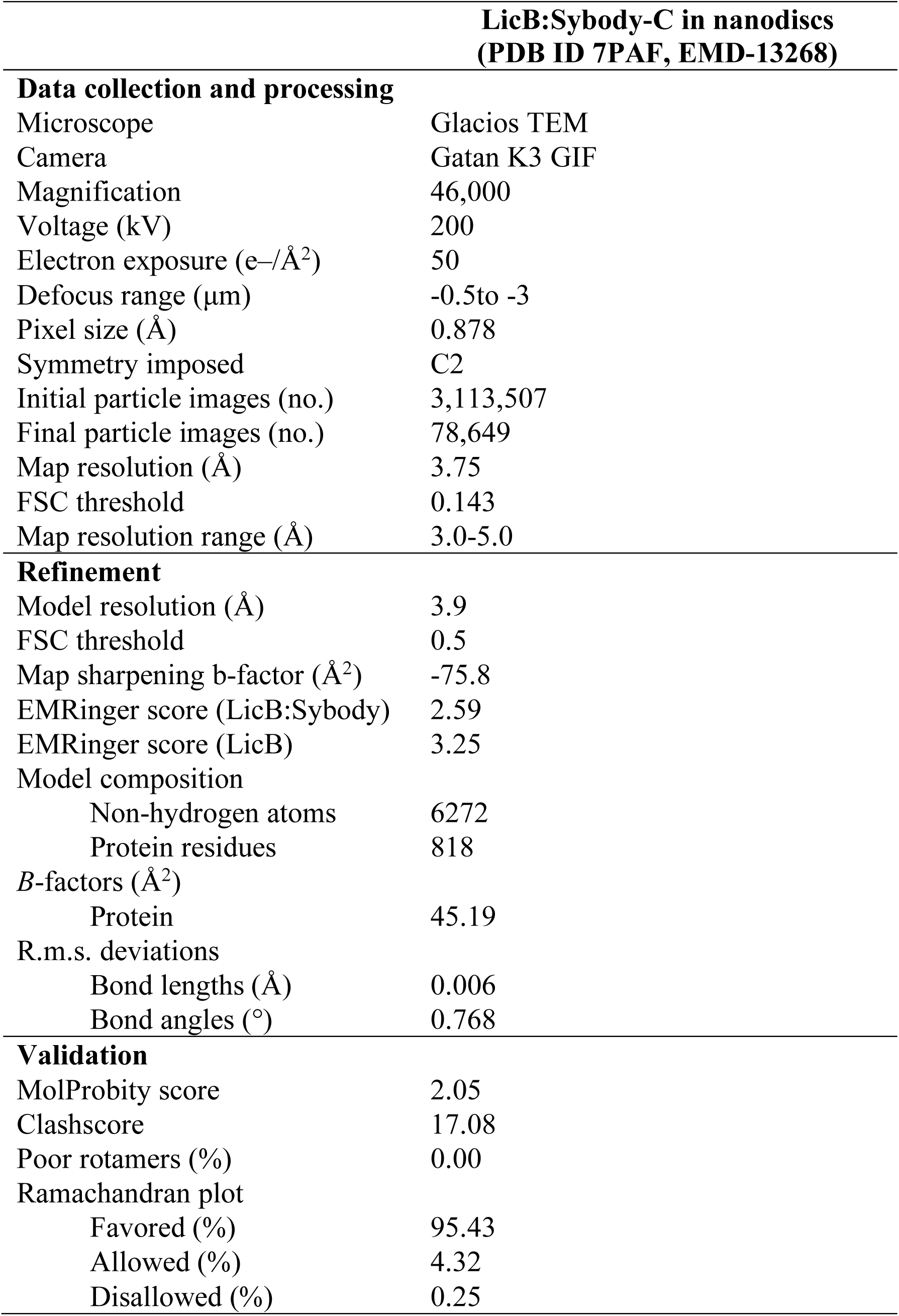
Cryo-EM data collection, refinement, and validation statistics

LicB displays a 10-TM helix topology with both N- and C-terminal domains facing the cytoplasm (**Fig. 3B,C**). LicB exhibits the DMT superfamily fold consisting of two ‘inverted’ structural repeats with antiparallel topology comprised by the N-terminal (TM1-TM5) and the C-terminal (TM6-TM10) domains (**Fig. 3B**). TM1–TM4 and TM6–TM9 form a central cavity that opens up towards the extracellular side of the membrane, indicating that the structure is in an outward-open conformation (**Fig. 3E**). Access to the central cavity from the cytoplasmic face is sealed off by multiple interactions between TM6-TM9 and TM7-TM8 and the cytoplasmic loops connecting TM6-7 and TM8-9. Multiple aromatic residues including W17 (TM1), Y109 (TM4), W167 (TM6), Y233 (TM9), Y236 (TM9), and Y255 (TM10) participate in formation of the central cavity (**Fig. 3C**). The structural fold exhibited by LicB is similar to that of other structurally characterized DMT transporters including the amino-acids exporter Yddg (*54*), the *Plasmodium falciparum* drug transporter PfCRT (*55*), the triosephosphate/phosphate antiporter TPT (*56*), and the SLC35 nucleotide sugar antiporters CST (*57, 58*) and Vrg4 (*59, 60*).

### Structure of choline-bound LicB in occluded state

A choline-bound structure, displaying a different conformational state to that observed by cryo-EM, was obtained after co-crystallization of *S. pneumoniae* LicB in presence of choline (**Fig. 4A, Fig. S10A,B and Table 2**). An important experimental aspect to solve this structure was to perform dehydration of LicB crystals, which improved diffraction resolution from 6 to 3.8 Å. Phases were determined by molecular replacement using a LicB protomer from the cryo-EM structure as search model. The overall electron density map was of good quality throughout the polypeptide chain, except for loops 2 and 7, which were not included in the final model (**Fig. 4A**). The crystal structure revealed a conformation occluded from both sides of the membrane (**Fig. 4B**). Access to the central cavity is sealed by extracellular interactions between helices TM1-TM4 and TM2-TM3, and the loops connecting TM1-2 and TM3-4, and by cytoplasmic interactions between TM6-TM9 and TM7-TM8 and the cytoplasmic loops connecting TM6-7 and TM8-9. A clear positive peak in the *Fo-Fc* map indicated the presence of choline in the central cavity (**Fig. 4A and Fig. S10B**). Residues coordinating the choline molecule include W17 (TM1), Y109 (TM4), W167 (TM6), Y233 (TM8), and Y255 (TM9), which form an ‘aromatic box’ surrounding the trimethylammonium group, whereas Y236 (TM8), and N252 (TM9) coordinate the hydroxyl end of the molecule. Computational docking confirmed that the same binding pocket could accommodate an acetylcholine molecule (**Fig. S11A**). The larger size of acetylcholine likely explains the lower affinity observed for this molecule (**Fig. 2B**). In agreement with the distribution of substrate binding residues described here for LicB, substrate-bound structures of other DMT transporters, including the triosephosphate/phosphate antiporter TPT (*56*) and the SLC35 nucleotide sugar antiporters CST (*57, 58*) and Vrg4 (*59, 60*), have shown that residues from TM1-TM4, from the first inverted repeat, and TM6-TM9, from the second repeat, are involved in substrate coordination. Thus, albeit the significant differences among substrates of DMT transporters, the arrangement of substrate binding residues seems to be conserved.

**Figure 4.**
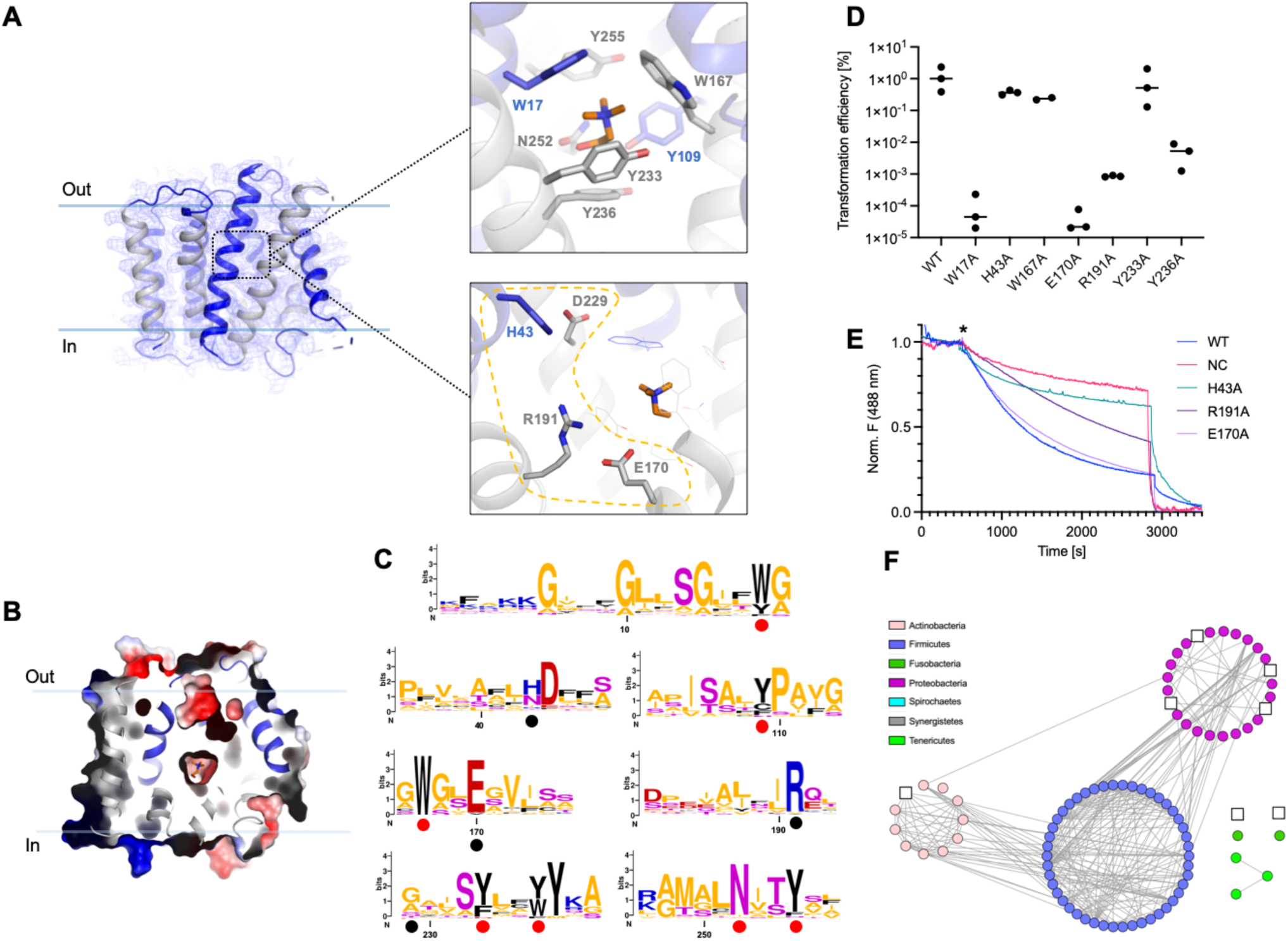
Crystal structure of choline-bound occluded *S. pneumoniae* LicB. **A**. LicB and *2Fo-Fc* electron density map contoured at 1.0 σ. Inverted repeats, TM1-5 and TM6-10, are indicated in blue and grey, respectively. (*Top*) Choline is shown with carbons colored in orange. (*Bottom*) Charged residues in the vicinity of the choline binding site. **B**. Surface electrostatic potential representation of LicB showing the central cavity closed to both sides of the membrane. Choline is shown in orange. **C**. Sequence logos of the regions containing the choline binding residues (red dots) and charged residues around the central cavity (black dots). Sequences are from LicB proteins analyzed in Fig. 1B (see Fig. S1). The y axis denotes positional information in bits. **D**. Transformation efficiency assays of *S. pneumoniae* D39V Δ*licB* cells complemented with WT *Plac-licB* or variants (n=3, biological replicates). **E**. Proton transport assay with WT LicB and variants in proteoliposomes. Representative time courses are shown (n=3, biological replicates). H^+^ influx was induced by establishing a membrane potential upon addition of the potassium ionophore valinomycin (star). The proton gradient was collapsed by the addition of CCCP at about 3000 seconds. NC indicates negative control (protein-free liposomes). **F**. Sequence similarity network as in Fig. 1B, showing bacteria species where residues E170 and R191 are not conserved (white squares, see Fig. S1).

**Table 2:** X-ray data collection and refinement statistics

| Data collection | LicB (PDB ID 7B0K) |
| --- | --- |
| Wavelength (Å) | 1.000031 |
| Space group | C 1 2 1 |
| Unit cell: |  |
| a/b/c (Å) | 128.90/43.44/126.82 |
| α/β/γ (°) | 90.0/120.43/90.0 |
| Resolution (Å) | 13-3.8 |
| Completeness (%) | 86.8(37.2) [98.9(97.2)] |
| No. measured reflections | 18226(690) [20849 (2305)] |
| No. unique reflections | 5593(196) [7167(694)] |
| I/σI | 4.7(2.3) [2.05(1.2)] |
| R-meas (%) | 14.6(67.7) [23.1(144.7)] |
| CC<sub>1/2</sub> (%) | 99.6(85.1) [99.5(66.3)] |
| **Refinement** |  |
| R<sub>work</sub>/R<sub>free</sub> (%) | 29.69/31.97 |
| R.m.s.d. Bonds (Å) | 0.002 |
| R.m.s.d. Angles (°) | 0.605 |
| Ramachandran plot: |  |
| Outliers (%) | 0.0 |
| Allowed (%) | 7.84 |
| Favored (%) | 92.16 |
| Average B-factor (Å<sup>2</sup>) | 107.0 |

Values in brackets are before anisotropic truncation Values in parentheses are for the last resolution shell $R_{merge} = \sum_{hkl} \sum_i |I_i(hkl) - \langle I(hkl) \rangle| / \sum_{hkl} \sum_i I_i(hkl)$

### Conserved choline binding residues are relevant for *S. pneumoniae* fitness

Sequence conservation analysis revealed that residues W17, Y109, W167, Y233, Y236, N252, and Y255, participating in coordination of choline at the central pocket are highly conserved among LicB proteins across different bacterial phyla (**Fig. 4C and Fig. S11B**), including the Gram-negative pathogen *H. influenzae* (**Fig. S11C**). To further show the importance of residues participating in choline binding, we performed complementation assays in *S. pneumoniae*. First, we constructed strains in which residues participating in formation of the ‘aromatic box’ surrounding the trimethylammonium group of choline (**Fig. 4A**) were exchanged to alanine (W17A, W167A, Y233A, Y236A) and the genes encoding these variants were cloned under the control of an IPTG-inducible promoter (P_*lac*_) and integrated at the ectopic ZIP locus as second copy of *licB* (*61*). Transformation assays were done aiming to replace the native *licB* gene with an erythromycin resistance cassette and transformants were plated with and without IPTG. As expected, wild type (WT) P_*lac*_*-licB* could complement the *licB* deletion (**Fig. 4D**). We obtained an average transformation efficiency of approximately 1% in the presence of IPTG while no colonies appeared in the absence of IPTG. Exchange to alanine of residues Y233 and W167 resulted in transformation efficiencies comparable to WT *licB*, whereas mutants W17 and Y236 resulted in significantly reduced transformation efficiencies indicating that these are essential residues for LicB function (**Fig. 4D**). Thus, slight modifications of the binding pocket of LicB are sufficient to perturb fitness of *S. pneumoniae*.

### LicB is a proton-dependent choline symporter

The central cavity of LicB displays multiple charged residues in close proximity to the choline binding site (**Fig. 4A**), with residues H43, E170, and R191, being conserved among LicB proteins from different bacterial species (**Fig. 4C and Fig. S11B**). The chloroquine resistant transporter PfCRT, which has been described to be a H^+^-coupled transporter, displays similar charged residues located at its central cavity (*55, 62*). Charged residues in central cavities of secondary transporters have been frequently associated with proton transport as these residues are prone to protonation and de-protonation due to fluctuations in their surrounding chemical environment during cycling. We wanted to test whether similarly to PfCRT, LicB could display H^+^-coupled activity. To test this, we performed membrane potential driven H^+^ transport assays with LicB reconstituted in proteoliposomes in presence of the fluorophore 9-amino-6-chloro-2-methoxyacridine (ACMA) (**Fig. 4E**). The robust fluorescence decrease observed upon the addition of valinomycin reflects H^+^ influx into LicB-WT proteoliposomes in contrast to the slower quenching of ACMA observed in protein-free liposomes (**Fig. 4E**). Similar experiments with variants E170A, R191A, and H43A reconstituted in proteoliposomes (**Fig. 4E**), revealed that H^+^ influx decreases for variants R191A and H43A, whereas E170A exhibited influx similar to that of WT LicB (**Fig. 4E**). Complementation assays in *S. pneumoniae* using the above-mentioned transformation assay, showed that H43A supports viability, whereas mutants E170A and R191A did not (**Fig. 4D**). Strikingly, E170 and R191 are particularly highly conserved residues among bacteria phyla where genes of the *lic* operon are found (**Fig. 4C**), with only very few exceptions as indicated in **Fig. 4F**. Taken together, these results led us to hypothesize that H^+^-coupled choline import is a highly conserved trait of LicB proteins across bacteria displaying phosphocholine decoration.

### Comparison of LicB outward-open and occluded conformations

Structures of other DMT transporters have been solved either in outward-open or occluded states, and so far, there are no structures available for inward-facing states (*54-56*). Modeling of different conformations have been possible thanks to the structural conservation of the DMT fold and the pseudo-symmetrical arrangement of TM helices in two inverted repeats (*38*). However, until now, LicB is the only DMT transporter with elucidated structures in two distinct structural states. Analysis of the conformational changes taking place during the transition from outward-open to occluded state, shows that TM3 and TM6 are the main players in closing the access to the central cavity as they move towards the core of the translocation pathway (**Fig. 5A,B**). During this transition, the extracellular halves of TM3 and TM6 tilt about 20°. An inward-facing model based on the pseudo-symmetrical arrangement of the two repeats, predict that the transition from occluded- to inward-open conformation would involve movements of the cytoplasmic halves of TM1 and TM8 (**Fig. 5B**). Large movements of the extracellular halves of TM3 and TM6 are favored by the flexibility introduced by glycine residues at their midsections. In TM3 this includes residues G83-G86; G168 in TM6; G14-G18-G20 in TM1; and G216-G220 in TM8. Similar flexible segments have been observed in YddG (*54*), TPT (*56*), Vrg4 (*60*), and CST (*57, 58*), and glycine residues at this same regions are present in LicB proteins, indicating that similar conformational changes are expected to occur in these proteins.

**Figure 5.**
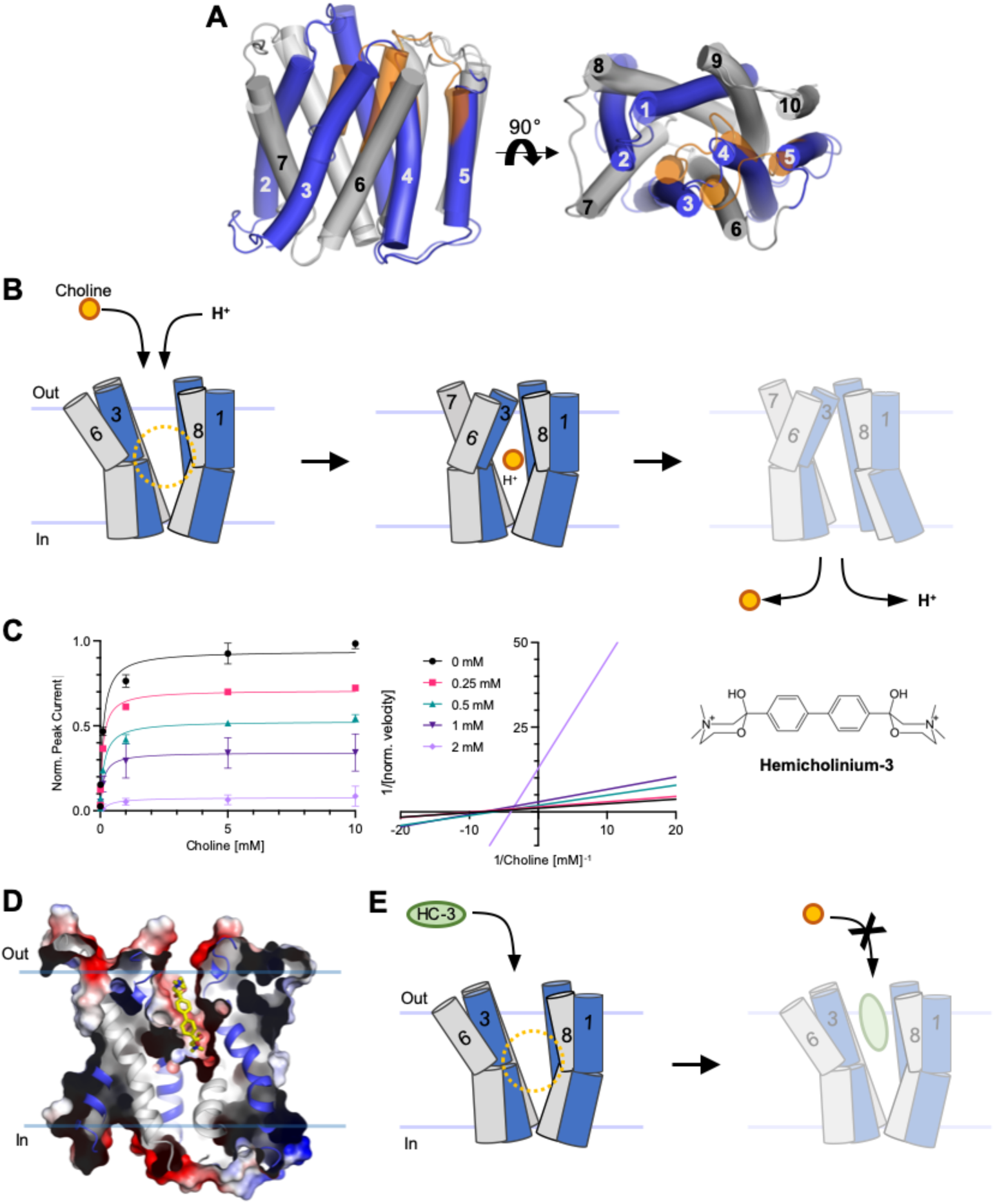
Alternating access model of choline import and inhibition by HC-3. **A**. Superposition of outward-open and occluded structures of LicB. Inverted repeats, TM1-5 and TM6-10, are indicated in blue and grey, respectively. Helices that undergo large conformational changes are shown in orange in the occluded state. **B**. Schematic of the proposed transport cycle depicting the movements of TM3 and TM6 that close the extracellular entry pathway, whereas the predicted movements of TM1 and TM8 open the cytoplasmic exit pathway. Choline (orange) and protons enter the binding cavity in the outward-open state. **C**. (*Left*) Inhibition of LicB by HC-3 measured by SSM-electrophysiology (K_i_=518 ± 31 µM). (*Right*) Lineweaver-Burk plot showing that HC-3 is a non-competitive inhibitor of LicB. Norm. velocity indicates normalization of maximum currents amplitude. **D**. Surface electrostatic potential representation of outward-open LicB showing a HC-3 molecule docked in the extracellular entry pathway. **E**. Schematic of the proposed model of inhibition by HC-3 (green). HC-3 blocks the extracellular entry pathway hindering the access of choline to the central binding site.

### LicB is inhibited by hemicholinium-3

The importance of LicB in maintaining the choline supply for phosphocholine synthesis, makes this transporter a very attractive target for drugs development. However, besides the inhibitory sybodies described here, there are no reported inhibitors of LicB. In particular, it would be important to develop small molecule inhibitors with the potential to cross the cell wall. Hemicholinium-3 (HC-3) is a well characterized inhibitor of the high-affinity choline transporter CHT1 (SLC5A7) (*63*), a member of the sodium-solute symporter family in which the structurally characterized members exhibit the conserved LeuT-fold (*64*). Albeit, the architecture of DMT and LeuT-fold transporters is markedly different, we wanted to test whether HC-3 could inhibit the choline transport activity of LicB. To test this, we performed SSM-electrophysiology measurements with LicB reconstituted in proteoliposomes, and evaluated electrogenic currents in presence of different concentrations of HC-3 (**Fig. 5C**). Our results show that at a concentration of 0.5 mM of HC-3, the electrogenic transport of choline is reduced about 50%, while at 2 mM, the activity is reduced about 90%. HC-3 displays non-competitive binding to LicB with a K_i_ of 518 ± 31 µM (**Fig. 5C**). This mode of inhibition indicates that HC-3 does not bind to the choline binding pocket, in agreement with the larger size of this molecule in comparison to choline. Computational docking of HC-3 to the outward-open structure of LicB revealed that it preferentially binds to the extracellular entrance pathway, blocking the access to the central binding site (**Fig. 5D,E**). Taking together, these results indicate that HC-3 is a genuine scaffold for future design of inhibitors targeting LicB proteins.

## Discussion

Phosphocholine synthesis is an essential part of the pathways that decorate TA and LPS in pathogenic bacteria. Considering the significant roles of these two biopolymers in adhesion to host cells, immune evasion, and bacterial persistence (*7, 8*), understanding the mechanisms of proteins involved in phosphocholine decoration is critical for the development of strategies aiming to counteract bacterial infections. Pathogens like *S. pneumoniae* and *H. influenzae* are unable to synthesize choline, thus, uptake of this molecule is a fundamental step in exploiting the advantages that TA or LPS phosphocholine decoration provide. The essential role of LicB in supplying with choline the protein machinery encoded by the *lic* operon to perform TA and LPS decoration, makes LicB and other proteins related to this process promising targets for drugs development. Thus, demonstrating how LicB works and elucidating architectural and mechanistic elements essential to its function is of significant relevance. The role of catalytic residues involved in choline binding and proton-coupling in *S. pneumoniae* LicB as described here, and their high conservation across divergent bacteria phyla, support the essential role of these mechanistic elements in supplying the phosphocholine synthesis pathway for decoration of TA and LPS. Indeed, a homology model of *H. influenzae* LicB reveals a central cavity with an ‘aromatic box’ and charged residues at the same positions as in *S. pneumoniae* LicB (**Fig. S11C**).

Despite the fact that inhibitory sybodies targeting LicB pose great potential to counteract bacterial pathogens, further strategies aiming to facilitate sybodies diffusion across their capsule and cell wall need to be developed. On the other hand, small molecule inhibitors, such as HC-3, are likely to inhibit LicB activity much more effectively under native conditions due to less restricted diffusion. Thus, the biochemical and structural characterization of the inhibitory mechanism of HC-3 set the basis for development of high specificity inhibitors.

The promiscuous selectivity of LicB is a striking finding that point towards an adaptation mechanism of *S. pneumoniae* under conditions with limited accessibility to free choline. We demonstrated that *S. pneumoniae* can catabolize acetylcholine and use the choline product for TA functionalization. However, the nature of the protein(s) that perform acetylcholine hydrolysis in *S. pneumoniae* and how conserved is this mechanism among other bacteria remains to be shown. Docking analysis of acetylcholine indicates that the highly conserved catalytic residues involved in choline binding, suffice to coordinate acetylcholine. Thus, we speculate that LicB proteins in other bacteria would be able to import acetylcholine as well.

From the structurally characterized DMT transporters, PfCRT is the most similar to LicB in terms of function as it performs symport towards the cytoplasm (*55, 62*). PfCRT exports drugs from the acidic digestive vacuole of intra-erythrocytic *P. falciparum* parasites (*62*). Comparison of outward-open LicB and outward-open PfCRT (PDB: 6UKJ), which superpose with an r.m.s.d. of 2.6 Å (Cα atoms), reveal similarities in the location of charged residues in their central cavities; H43_LicB_/H97_PfCRT_, E170_LicB_/D137_PfCRT_, R191_LicB_/R231_PfCRT_, and D229_LicB_/D326_PfCRT_. Like LicB, PfCRT also makes use of the electrochemical proton gradient (*65*). Proton-coupled transport by LicB implicates that acidification of the external media would result in faster choline uptake. This might be relevant at the natural niche of pathogens like *S. pneumoniae* and *H. influenzae*, the human nasopharynx, which displays mild acidic conditions (5.0 < pH < 6.5) (*66, 67*). In the absence of a proton gradient, neutral pH, or in the presence of a mutation that disrupts proton transport (H43A), transport of choline is driven by the membrane potential and likely by a choline concentration gradient since the phosphocholine synthesis pathway quickly diminishes the concentration of free choline in the cytoplasm (*36, 68, 69*).

The cryo-EM structure of LicB in nanodiscs revealed a homodimer arrangement. Dimerization has been observed in crystal structures of DMT transporters including the triosephosphate/phosphate antiporter TPT (*56*), the GDP-mannose transporter Vrg4 (*59, 60*) and the CMP-sialic acid transporter CST (*57, 58*). Strikingly, the dimer arrangement displayed by LicB is very different to that observed in the crystal structures of these DMT transporters. While the LicB interprotomer interactions involve TM1, TM8, and TM9 (**Fig. 3D**), the interactions observed in the structures of TPT, Vrg4, and CST, involve TM5 and TM10. Interestingly, the packing of the crystal structure of LicB shows a symmetry partner that interact through TM5 and TM10, thus, displaying the same dimer topology as that described for TPT, Vrg4, and CST (**Fig. S10C**). Although for TPT, it is not known whether the dimer is functionally relevant, it has been shown that dimerization of Vrg4 and CST allows faster transport and enhances protein stability (*57, 59*). Since LicB dimerization was observed during purification, albeit in low proportion (**Fig. S12**), and lipid nanodiscs represent a native-like environment due to the presence of a lipid bilayer and absence of detergent micelles, the arrangement of LicB molecules in the cryo-EM structure might represent a functional dimer, but this remains to be shown.

More broadly, we showed a comprehensive mechanistic analysis of a DMT transporter essential for bacterial cell wall modification. Due to the widespread importance of phosphocholine decoration of teichoic acids and lipopolysaccharide in bacterial pathogens, our results constitute a significant advance that is likely to contribute to the development of new strategies to modulate interactions with the host. Indeed, sybody-like molecules or derivatives of HC-3 represent promising chemical structures that will facilitate the engineering of new molecules targeting LicB proteins.

## Materials and Methods

### LicB expression, purification and nanodiscs reconstitution

The gene encoding *S. pneumoniae* LicB with an N-terminal 10×His affinity tag in a modified pET-19b vector (Novagen) was overexpressed in *E. coli* BL21-Gold (DE3) (Stratagene) cells. The cells were grown at 37°C in Terrific Broth medium supplemented with 1% glucose (w/v) and 100 µg/ml ampicillin. Protein overexpression was induced with 0.2 mM β-D-1-thiogalactopyranoside (IPTG) and cells were incubated for 1 hour before harvesting. Frozen cell pellets were resuspended in 50 mM Tris-HCl, pH 8.0; 500 mM NaCl; 5 mM β-mercaptoethanol and 0.5 mM PMSF. Cells were disrupted, the membrane fraction was separated by ultracentrifugation and subsequently flash frozen with liquid nitrogen until further use. Frozen membranes were solubilized for 2 hours at 4°C in 50 mM Tris-HCl, pH 8.0; 200 mM NaCl; 15 % glycerol (v/v); 20 mM imidazole, pH 8.0; 1% Lauryl Maltose Neopentyl Glycol (w/v) (LMNG, Anatrace); and 2 mM β-mercaptoethanol. After centrifugation the supernatant was loaded onto a Ni-NTA superflow affinity column (Qiagen), pre-equilibrated on 50 mM Tris-HCl, pH 8.0; 500 mM NaCl; 10 % glycerol; 20 mM imidazole, pH 8.0; 0.2% LMNG and 2 mM β-mercaptoethanol and subsequently washed with buffer containing 50 mM Tris-HCl, pH 8.0; 500 mM NaCl; 10 % glycerol (v/v); 50 mM imidazole, pH 8.0; 0.2% LMNG and 2 mM β-mercaptoethanol. The protein was eluted with buffer containing 50 mM Tris-HCl, pH 8.0; 200 mM NaCl; 10 % glycerol (v/v); 500 mM imidazole, pH 8.0; 0.2% LMNG and 2 mM β-mercaptoethanol. The buffer was exchanged to 10 mM Tris-HCl, pH 8.0; 150 mM NaCl and 0.012% LMNG with PD-10 columns (GE healthcare) for the removal of imidazole. The 10×His affinity tag was removed by overnight incubation with Tobacco Etch Virus (TEV) protease, which was later removed by passing through a Ni-NTA affinity column. LicB was further purified by size exclusion chromatography (SEC) with running buffer 10 mM Tris-HCl, pH 8.0; 150 mM NaCl and 0.012% LMNG, using a Superdex 200 Increase 10/300 GL column (GE Healthcare). Purified LicB was reconstituted in MSP1D1 nanodiscs using a ratio of 3:9:7:175 (LicB:Sybody:MSP1D1:lipids) in a buffer containing 50 mM Tris-HCl, pH 8.0; 50 mM NaCl and 10 % glycerol (v/v). The lipid mixture used consist of 16:0-18:1 POPG:*E. coli* polar lipid extract (Avanti) in a 3:1 (w:w) ratio. Detergent was removed by adding Bio-beads SM2 (BioRad). After Bio-beads removal the mixture was centrifuged and loaded on a Superdex 200 Increase 10/300 GL (GE Healthcare) column equilibrated with buffer 50 mM Tris-HCl, pH 8.0; 50 mM NaCl. The peak corresponding to LicB:Sybody in nanodiscs was collected and used for single particle cryo-EM studies.

### Synthetic nanobody (sybody) selection

LicB carrying an N-terminal Avi-tag, followed by a SSGTSS linker sequence to warrant efficient biotinylation, was expressed and purified as described above. Enzymatic biotinylation was performed using recombinant BirA in presence of 0.5 mM biotin, 20 mM Magnesium acetate, 20 mM ATP, and 5% glycerol(*70*). The reaction was incubated for 16 hours at 4°C. Protein biotinylation was confirmed by SDS-PAGE after incubation of the reaction product with Streptavidin. Biotinylated LicB was desalted with buffer 10mM Tris-HCl pH 8.0, 150 mM NaCl, 0.012% LMNG, and His-tagged BirA was removed using a Ni-NTA superflow affinity column. The protein was aliquoted at a concentration of 5 μM, flash frozen in liquid nitrogen, and stored at -80°C until used for nanobodies selection. Sybody selection was performed as previously described(*46*). In short, sybodies were selected against biotinylated LicB in presence of 5 mM choline using the three sybody libraries concave, loop and convex. After one round of ribosome display, two rounds of phage display were performed, switching the panning surface in each round. During the second round of phage display, an off-rate selection was carried using non-biotinylated LicB at a concentration of 5 µM. The enrichment was monitored throughout the selection process by qPCR, which looked ideal with values of 4.6 (concave), 2.5 (loop) and 3.3 (convex) after the first round of phage display and 1922 (concave), 107 (loop) and 70 (convex) after the second round of phage display. 95 single clones of each library were screened by ELISA resulting in 40 hits. Sanger sequencing revealed 29 unique sybodies, of which 26 showed a monodisperse peak at the expected elution volume on a Sepax SRT-10C SEC-100 column. From these, the best behaved sybodies during expression and purification (12 sybodies, all belonging to the convex sybody library, **Fig. S4**) were selected for further functional characterization.

### Sybody expression and purification

Sybodies were expressed and purified as previously described(*46, 47*), with minor modifications. In brief, the sybody encoding gene in the pSBinit vector (Addgene #110100) was transformed into *E. coli* MC1061 competent cells, which were then grown at 37°C in Terrific broth media supplemented with 0.004 % glycerol, 100 µg/ml ampicillin and 100 µg/ml streptomycin. Overexpression of the protein was induced with 0.02% L-arabinose at an OD_600_ of 0.7, followed by 15 hours incubation at 22°C before harvesting. The cell pellet was resuspended in 40 mM Tris-HCl, pH 7.4; 500 mM NaCl; 0.5 mM PMSF and cells were lysed using a tip sonicator. The cell debris was discarded by centrifugation and the supernatant was loaded on a Ni-NTA affinity chromatography column equilibrated with 40 mM Tris-HCl, pH 7.4; 150 mM NaCl; 50 mM imidazole. The column was washed with 40 mM Tris-HCl, pH 7.4; 150 mM NaCl; 50 mM imidazole, and the protein eluted with 40 mM Tris-HCl, pH 7.4; 150 mM NaCl; 300 mM imidazole. Imidazole was then removed using a PD-10 column (GE Healthcare). The concentrations of sybody preparations were determined by measuring A_280_ and the quality of the purified sybodies was assessed by SDS-PAGE and SEC.

### Sybody binding constant determination

Kinetic characterization of sybodies binding to biotinylated LicB was performed using grating-coupled interferometry (GCI) on a WAVEsystem instrument (Creoptix AG). Biotinylated LicB was captured onto a Streptavidin PCP-STA WAVEchip (polycarboxylate quasi-planar surface; Creoptix AG) to a density of 1700 pg/mm^2^. Sybodies were injected at increasing concentrations ranging from 5 nM to 405 nM using a three-fold serial dilution and 5 different concentrations in buffer 10 mM Tris-HCl pH 8.0, 150 mM NaCl, 20 mM imidazole supplemented with 0.02 % LMNG with or without 5 mM choline. Sybodies were injected for 200 s at a flow rate of 50 µl/min. Dissociation recording was set to 900 s to allow the return to baseline. Blanks were injected after every second analyte injection. All sensorgrams were recorded at 25°C and the data was analyzed on the WAVEcontrol software (Creoptix AG). Data were double-referenced by subtracting the signals of buffer injections (blanks) and by subtracting the signals of a reference channel. A Langmuir 1:1 model was used for data fitting. For Sybody-C the two highest concentrations were excluded from data fitting, due to unspecific interaction with the flow channels at high concentrations.

### Sample preparation and cryo-EM data acquisition

A sample of nanodiscs reconstituted LicB:sybody-C complex was concentrated between 1 to 1.5 mg/ml using a Vivaspin concentrator with a 30,000 Da cutoff (GE Healthcare). Cryogenic samples were prepared using a Mark IV Vitrobot (Thermo Fisher) at 95% humidity at 4°C. 5 µl of the sample was applied to glow discharged Quantifoil R1.2/1.3 300-mesh copper holey carbon grids, blotted for 4 seconds using a blotting force of 3. The grids were flash frozen in a mixture of propane and ethane, then cooled with liquid nitrogen. Movies were recorded with SerialEM on a Glacios microscope (Thermo Fisher) operated at 200 kV, equipped with a K3 direct electron detector (Gatan). Images were recorded with a defocus range of 0.5 and 3 µm and a pixel size of 0.878 Å/pixel at a nominal magnification of 46,000 ×. Each micrograph was dose-fractionated to 25 frames under a dose rate of 12.5 electrons per pixel per second, with a total exposure time of 4 seconds, resulting in a total dose of about 50 e^−^/Å^2^.

### Cryo-EM data processing

Data processing was carried out entirely in cryoSPARC v3.2.0(*71*). Beam-induced drift of 8,229 raw movies was corrected and the movies aligned using patch motion correction. The contrast transfer function (CTF) of each aligned micrograph was determined using patch CTF estimation. Micrographs were Fourier-cropped once during patch motion correction to adjust the pixel size to 0.878 Å per pixel. Micrographs were classified and some discarded, based on CTF fit resolution, relative ice thickness, and total full-frame motion, resulting in 8,008 micrographs. A small subset of micrographs was used to adjust parameters for optimal picking of particles, which was performed using blob picker. 3,113,507 particles were extracted with a box size of 384 pixel and Fourier-cropped to 192 pixels, resulting in a pixel size of 1.756 Å per pixel. Multiple rounds of 2D classification with a batch size of 200 particles per class were performed to discard bad particles. The selected particles were used to perform sequential *ab initio* reconstructions with 3 or 4 classes, no similarity, and C1 symmetry. The best-resolved class was used as reference for multiple rounds of heterogeneous refinement with the particle sets selected from the preceding round of 3D classification. After heterogeneous refinement, 148,064 good particles were re-extracted, resulting in a pixel size of 0.878 Å per pixel, and further refined in a non-uniform (NU) refinement(*71*) using C2 symmetry, input reconstruction map filtered to 12 Å, and per-group CTF parameters. NU-refinement resulted in a map with a 4.2 Å resolution, according to the 0.143 cut-off criterion(*72*). This reconstruction map was used to generate templates for automated template-based picking from all good micrographs. Particles were re-extracted and a similar processing procedure as that described above was applied. After NU-refinement, a set of 78,649 particles were locally CTF refined before local resolution refinement, which resulted in a map with a 3.75 Å resolution, according to the 0.143 cut-off criterion. The local resolution was estimated with cryoSPARC v3.2.0(*71*) and the directional resolution estimation was done using the 3DFSC server(*73*).

### Model building

Model building was performed in Coot(*74*). The cryo-EM density of the LicB dimer was of sufficiently high resolution to unambiguously build LicB polypeptide chain into the cryo-EM density. The sybody was only partially resolved, with the part interacting with LicB being its best resolved region. Thus, a homology model of the sybody (based on PDB: 5m14) was positioned by rigid body in the density map, accompanied of manual building of the CDR3 region. Models were improved by iterative cycles of real-space refinement in PHENIX(*75*) with application of secondary structure constrains. Validations of models were performed in PHENIX. The final refined structure has 95.43% of residues in the Ramachandran favored region; 4.32% in Ramachandran allowed; and 0.25% as Ramachandran outliers. Figures of models and densities were made in PyMol (The PyMOL, Molecular Graphics Systems, Schödinger, LLC), and ChimeraX(*76, 77*).

### LicB crystallization

LicB in buffer 10 mM Tris-HCl, pH 8.0; 150 mM NaCl; 0.012% LMNG was concentrated up to 8 mg/ml using a 30 kDa MWCO Vivaspin 20 concentrator (GE healthcare). Extensive optimisation of crystal conditions using sitting-drop and hanging-drop vapour diffusion and other post-crystallisation treatments yielded plate shaped crystals after 3-4 days of incubation at 20°C. The reservoir conditions contained 1.5 mM choline; 100 mM HEPES, pH 7.5; and 33% PEG 400. Optimised crystals were dehydrated and cryoprotected by gently increasing PEG 400 concentration in the drop. Crystals were scooped from the drop at different time points followed by flash freezing by immersion in liquid nitrogen.

### Data collection

LicB crystals diffracted X-rays up to about 6-7 Å resolution in general. Drop dehydration through air exposure over 30 minutes led to crystals that diffracted X-rays to higher resolution(*78*). A data-set collected from one LicB crystal that showed anisotropic diffraction and belonged to space group C121 was used to determine the structure. The unit cell constants of this crystal were a = 128.9 Å, b = 43.44 Å, c = 126.82 Å and α = 90º, β = 120.43º, γ = 90º (**Table 2**). Data was processed and merged with XDS(*79, 80*) and anisotropic scaling/ellipsoid truncation was performed(*81*). Resolution limits after ellipsoid truncation were a* = 4.0 Å, b* = 3.5 Å, and c* = 4.2 Å. Karplus CC* (Pearson correlation coefficient) based data cutoff approach was used to determine the usable resolution of the data-sets(*82*). The resolution limit was set considering a CC_1/2_ > ∼40% based on data merging statistics and a CC* analysis against unmerged intensities in Phenix package(*75*) satisfying Karplus CC* against CC_work_ and CC_free_ criteria, as well as, R_free_ of the highest resolution shell against the refined structure being less than or equal to ∼50%. A second criterion for limiting the resolution was the overall completeness percentage observed after anisotropic ellipsoid truncation, which was kept above 85%. Diffraction data was collected at the beamline X06SA at the Swiss Light Source (SLS, Villigen).

### LicB crystal structure determination

The crystal structure of LicB was solved by molecular replacement using the program PHASER(*83*). The model generated by single particle cryo-EM was used as reference. Multiple rounds of refinement in Phenix(*75*) and model building using Coot(*74*) were performed. Map sharpening was used to facilitate model building. X-ray data and refinement statistics are given in **Table 2**. The final refined structure has R_work_ = 29.69% and R_free_ = 31.97%, with 92.16% of residues in the Ramachandran favored region; 7.84% in Ramachandran allowed; and 0.0% as Ramachandran outliers. Molecular graphics were created in PyMOL (The PyMOL, Molecular Graphics Systems, Schödinger, LLC). Surface electrostatics were calculated with the APBS PyMOL plugin.

### Reconstitution in proteoliposomes

Lipid mixtures of POPE:POPG 3:1 or ECPL:PC 3:1 (w:w) (Avanti), were mixed, dried from chloroform, and solubilised in buffer 20 mM Tris, pH 8.0; 150 mM NaCl; 2 mM β-mercaptoethanol, to a final concentration of 20 mg/ml. The lipids were flash frozen with liquid nitrogen and stored at -80°C. Unilamellar liposomes were formed after a 1:1 dilution in buffer and extrusion through a polycarbonate filter (400-nm pore size). After addition of 0.2% DDM, liposomes were mixed with LicB to have a 1:50 TM protein:lipid ratio (w:w). Detergent was removed with Bio-beads SM2 (BioRad). Proteoliposomes at a final concentration of 20 mg/ml of lipids were resuspended in 10 mM HEPES, pH 7.3; 100 mM KCl; 5 mM choline for the proton transport assay, and in 20 mM Tris, pH 8.0; 150 mM KCl; 2 mM β-mercaptoethanol for the SSM-based electrophysiology assays. The proteoliposomes were aliquoted, flash frozen in liquid nitrogen and stored at -80°C until further use.

### SSM-based electrophysiology

SSM-based electrophysiology was conducted using a SURFE2R N1 instrument (Nanion Technologies) following published protocols(*48, 84*). 3 mm SURFE2R N1 single sensors (Nanion Technologies) were alkylated by adding 100 µl of 0.5 mM thiol solution (1-octadecanethiol in isopropanol), subsequently incubating for 1 hour at room temperature in a closed petri dish. The sensor was then washed with isopropanol and miliQ water. 1.5 µl of lipid solution (7.5 <u>µg/µl</u> 1,2-diphytanoyl-sn-glycero-3-phosphocholin in n-decane) was applied on the surface of the gold electrode followed by immediate addition of 100 µl of SSM-buffer (50 mM Tris-HCl, pH 8.0; 5 mM MgCl_2_; 150 mM KCl). The proteoliposome suspension at a 20 mg/ml lipid concentration was diluted 1:20 in SSM-buffer and the mixture was sonicated for 20 seconds. 10 µl of the diluted proteoliposomes were added to the chip before centrifugation at 2,000 × g for 30 minutes. The conductance and capacitance were measured to ensure the quality of the SSM before each measurement. A conductance value below 5 nS, and capacitance between 15 - 35 nF were considered acceptable. Activating buffers (buffer-A) were prepared from a large stock of non-activating buffer (buffer-B: SSM-Buffer) by the addition of the substrate choline or other compounds such as arsenocholine, and acetylcholine at different concentrations ranging between 1 µM to 30 mM. Peak current values were determined for each individual substrate at different concentrations and plotted to calculate EC_50_ values. For the determination of the inhibitory constants of hemicholinium-3 (HC-3), the choline concentration was adjusted to 0.5, 1, 5, 10 mM choline in buffer-A, respectively and measured in the presence of 0.25, 0.5, 1 and 5 mM HC-3 present in buffer-A and buffer-B. Assays with sybodies were performed at a constant concentration of choline at 5 mM in buffer-A and 500 nM sybody in buffer-A and buffer-B as previously described(*48*). Prior to the measurements in presence of sybody, the maximal peak current was determined with 5 mM choline in buffer-A and used to normalise the peak currents in presence of sybody. Data for each sybody was collected from 3 independent sensor preparations and measured in triplicates. Analysis of the data was performed in OriginPro (OriginLab Corporation) and GraphPad Prism 9 (GraphPad Software).

### Protons transport assay

POPE:POPG (3:1) proteoliposomes were thawed and 20 µl of the proteoliposomes were sonicated and diluted to 500 µl with buffer containing 10 mM HEPES, pH 7.3, 10 mM KCl; 90 mM NaCl; 5 mM choline, and 0.75 µM 9-amino-6-chloro-2-methoxycridine (ACMA), before the assay. Time course fluorescence was measured at 20°C using a Jasco Fluorimeter. The change in ACMA fluorescence was detected using an excitation wavelength of 410 nm and an emission wavelength of 480 nm. After equilibration of the system, H^+^ and choline influx was initiated by establishing a membrane potential by the addition of the K^+^-selective ionophore valinomycin (5 nM). The activity was assessed for 1,200 seconds. The proton gradient was collapsed by the addition of 0.5 µM carbonyl cyanide m-chlorophenyl hydrazone (CCCP). Time courses were repeated at least three times for each individual experiment.

### Size-exclusion chromatography coupled to multi-angle light scattering

SEC-MALS measurements of LicB were performed at 18°C in 10 mM Tris-HCl, pH 8.0; 150 NaCl; 0.012% LMNG using a GE Healthcare Superdex 200 Increase 10/300 GL column on an Agilent 1260 high performance liquid chromatography. The column was equilibrated overnight for a stable baseline before data collection. Monitoring of the elution was carried out with a multi wavelength absorbance detector at 280 nm and 254 nm, the Wyatt Heleos II 8+ multiangle light-scattering detector, and a Wyatt Optilab rEX differential refractive index detector. Inter-detector delay volumes, light-scattering detector normalization and broadening corrections were calibrated using a 2 mg/ml bovine serum albumin solution (ThermoPierce) and standard protocols in ASTRA 6 (Wyatt Technologies). Weight-averaged molar mass, elution concentration, and mass distribution of the samples were calculated using the ASTRA 6 software (Wyatt Technologies). The dn/dc for the detergent LMNG was assumed to be 0.146 mg/ml according to experimental data(*85*).

### Mutagenesis

Point mutations H43A and R191A were introduced by site-directed mutagenesis using primers(*86*) (**Table S1**). Point mutations Y233A, Y236A, N252A, W17A, W167A and E170A were introduced by cloning using gBlock gene fragments (Integrated DNA Technologies) (**Table S1**).

### Sequence-similarity network

A sequence-similarity network of LicB proteins from multiple bacteria species (nodes) was generated using the EFI-EST webserver(*87*). Bacteria species were selected considering that *lic* operon genes encoding LicB and proteins that perform choline activation (LicA and LicC) are present(*88*). Sequences wre visualized with 40% identity and organized by phylum in Cytoscape(*89*). The phylogenetic tree showing the evolutionary relationships among the selected bacteria species was generated with iTOL(*90*).

### Structural sequence-conservation analysis

The sequence conservation analysis shown in **Fig. 10B** was computed using the ConSurf server(*91*). Briefly, LicB homologues displaying at least 35% identity were selected from a protein sequence BLAST search on the NCBI database using *S. pneumoniae* LicB protein sequence as a query. We then generated a multiple-sequence alignment using the HHMER algorithm provided by ConSurf, with conservation scores plotted in PyMOL (The PyMOL, Molecular Graphics Systems, Schödinger, LLC).

### Docking of hemicholinium-3 and acetylcholine

Docking of hemicholinium-3 and acetylcholine to the LicB structures was done with Autodock Vina(*92*). These molecules were downloaded in the SDF format from the ZINC database(*93*) and converted into a PDBQT format using Open Babel(*94*). Docking was carried out over a search space of 28 Å × 56 Å × 30 Å in the outward open state and 38 Å × 34 Å × 42 Å covering the entire entry pathway to the central cavity.

### Construction of mutants in *S. pneumoniae*

Strain VL4243 (*S. pneumoniae* D39V, prsI::PF6-lacI-tetR (gen), bgaA::Plac-licB (tet)) was made as follows. A second copy of *licB* under control of the IPTG-inducible P_*lac*_ promoter(*95*) was integrated in the genome of parental strain D39V at the *bgaA* locus. Golden Gate assembly of three parts was used. First part is the amplification by PCR using chromosomal DNA of strain VL1998 (lab collection) as template with primers OVL5139 and OVL5623 (**Table S1**). The resulting product contained the upstream homologous region to integrate at the *bgaA* locus, the tetracycline marker (*tet*), P_*lac*_ and the RBS. The second part is the amplification by PCR of *licB*, without its own RBS, from D39V with primers OVL5624 and OVL5625. The third part is the amplification by PCR using chromosomal DNA of strain VL1998 as template with primers OVL5626 and OVL2082 resulting on the downstream homologous region to integrate at the *bgaA* locus. After purification, the three parts were digested and ligated together during an assembly reaction with T4 DNA ligase buffer (Vazyme), T4 DNA ligase (Vazyme) and Eps3I (NEB). The assembly mixture was incubated in a thermocycler (PCR Max) by cycling 25 × the series 1.5 minutes at 37°C and 3 minutes at 16°C. The sample was then incubated for 5 minutes at 37°C and subsequently incubated for 10 minutes at 80°C. Strain VL333 (prsI::PF6-lacI-tetR (gen)(*96*)) was transformed with the assembly mixture and transformants were selected using tetracycline (µg/ml). The *bgaA* locus of the resulting strain, VL4243, was confirmed by sequencing. To create seven different point mutations in *licB*, the desired amino acids were changed for alanine in the second copy of *licB* at *bgaA*. Golden Gate assembly was used to amplify two parts per point mutation. The first part is the amplification by PCR of the upstream region of *bgaA::Plac-licB* from VL4243 until the desired point mutation with forward primer OVL2077 and reverse primer for each point mutation, OVL6035 (Y233A), OVL6037 (Y236A), OVL6039 (W17A), OVL6041 (W167A), OVL6043 (E170A), OVL6045 (R191A) or OVL6047 (H43A). The second part is the amplification by PCR of *bgaA::Plac-licB* from VL4243, started at the point mutation location until the downstream homologous region to integrate at the *bgaA* locus. Different forward primers were used for each point mutation, OVL6036 (Y233A), OVL6038 (Y236A), OVL6040 (W17A), OVL6042 (W167A), OVL6044 (E170A), OVL6046 (R191A), OVL6048 (H43A), the reverse primer was identical, OVL2082. After purification, the two parts were digested with Esp3I and ligated together during an assembly reaction with T4 DNA ligase buffer (Vazyme), T4 DNA ligase (Vazyme) and Eps3I (NEB). Strain VL333 was transformed with the assembly mixture with tetracycline selection. The *bgaA* region containing the P_lac_-*licB* genes of the resulting strains, VL4250 (Y233A), VL4251 (Y236A), VL4252 (W17A), VL4253 (W167A), VL4254 (E170A), VL4255 (R191A), VL4256 (H43A) were confirmed by sequencing.

### Transformation efficiency assay

Transformation efficiencies were determined to test whether mutated *licB* alleles would support growth as only copy of *licB* in the pneumococcal genome. Transformation assays were performed by transforming DNA of a *licB::ery* cassette to the strains with both copies of *licB* (without and with point mutation). DNA of *licB::ery* was obtained by amplification by PCR of strain VL4249 (lab collection) with primers OVL5635 and OVL5636. To transform *S. pneumoniae*, cells were grown in C+Y medium (pH 6.8) at 37°C to an OD of 0.1 at 595 nm. Subsequently, cells were treated for 12 minutes at 37°C with synthetic CSP-1 (100 ng /ml) and incubated for 20 minutes at 30°C with the transforming DNA (*licB::ery*). After incubation with the transforming DNA, cells were grown in C+Y medium (pH 6.8) at 37°C for 60 minutes. *S. pneumoniae* transformants were selected by plating, in triplicates, inside Columbia agar supplemented with 4% of defibrinated sheep blood (Thermo Scientific Oxoid) with 0.5 µg/ml erythromycin and 0 or 0.1 mM IPTG. To obtain the viable count, the transformant mix was diluted 1,000 or 10,000 times and plated without induction and selection, in triplicate, inside Columbia agar supplemented with 4% of defibrinated sheep blood. Transformation efficiency in percentage is calculated as the number of transformants divided by the number of total CFUs (viable count).

### *S. pneumoniae* sybodies susceptibility assays

Growth assays of unencapsulated *S. pneumoniae* strain VL567 (*cps::chl*) were performed in presence or absence of sybodies-A, -B, -C and -D at three different concentrations. Bacteria were grown in microtiter plates (CytoOne, CC 7672-7596) and incubated at 37°C, without shaking, in a Tecan i-Control infinite 200 PRO. Bacterial growth was monitored by measuring the optical density at 595 nm every 10 minutes during 8 hours. Measurements were performed using three replicates per condition. Strain VL567 was grown in C+Y (pH 6.8) until the optical density at 595 nm reached 0.1. Cells were spun down to remove the media and replaced by fresh C+Y medium. These washed pre-cultures were diluted 20-fold in C+Y without sybody or in presence of the four sybodies at different final concentrations per well.

### *S. pneumoniae* teichoic acids extraction and analysis

Cells were grown in 5 ml modified C+Y media(*97*) without choline or yeast containing radiolabeled [^3^H]-acetylcholine either at the acetyl group or the amino-group at 37°C until the OD reached 0.5 using 600 nm. The cell solution was split in two equal halves to LTA and WTA extraction. For LTA extraction cells were centrifuged at 5,000 × g for 5 minutes and the pellet washed twice with 1 ml buffer 20 mM MES, pH 6.5; 0.5 M sucrose; 20 mM MgCl2, before resuspending it in 1 ml of the same buffer. Proteoplasts were generated by the addition of 20 mg/ml lysozyme and 100 units mutanolysin. The mixture was incubated for 30 minutes at 37°C. The protoplasts were pelleted at 5,000 × g for 5 minutes and lysed in ice cold buffer containing 20 mM HEPES, pH 8.0; 100 mM NaCl; 1 mM DTT; 1 mM MgCl_2_; 1 mM CaCl_2_; 2 × complete protease inhibitors; 6 μg/ml RNAse A; 6 μg/ml DNAse I. Unbroken spheroplasts were removed by centrifugation at 5,000 × g for 10 minutes. The membrane fraction was collected by ultracentrifugation for 1 hour at 100,000 × g and at 4°C. The pellet was resuspended in 100 µl of buffer containing 50 mM MES, pH 6.5; 150 mM NaCl and the solution was flash frozen and stored at -80°C for further use. For WTA extraction cells were pelleted by centrifugation at 5,000 × g for 5 minutes and resuspended in 1 ml buffer containing 50 mM MES, pH 6.5, before a second centrifugation at 5,000 × g for 5 minutes and resuspension in 1 ml buffer containing 50 mM MES, pH 6.5; 4% (w/v) SDS. The solution was incubated at 100°C for 1 hour. The sacculi were collected with a 5,000 × g centrifugation for 5 minutes and washed with 1 ml buffer containing 50 mM MES, pH 6.5. The sample was centrifuged in a clean tube at 14,000 × g for 5 minutes. The pellet was washed with 1 ml buffer containing 50 mM MES, pH 6.5; 4% (w/v) SDS, twice with 1 ml buffer containing 50 mM MES, pH 6.5; 2% (w/v) NaCl and one in buffer containing 50 mM MES, pH 6.5. The sample was centrifuged at 14,000 × g for 5 minutes, resuspended in 1 ml buffer containing 20 mM Tris-HCl, pH 8.0; 0.5% (w/v) SDS; 20 µg of proteinase K. The sample was incubated for 4 hours at 50°C while shaking at 1,000 rpm. The pellet was collected by centrifugation at 14,000 × g for 5 minutes and washed with 1 ml buffer containing 50 mM MES, pH 6.5; 2% (w/v) NaCl. The sample was then washed three times with distilled water. The pellet was collected by centrifugation and hydrolyzed in 0.5 ml alkaline solution of 1 N sodium hydroxide. The mixture was incubated for 16 hours at 25°C while shaking at 1,000 rpm. Insoluble material was pelleted by centrifugation at 14,0000 × g for 5 minutes and the supernatant transferred into a clean tube. 125 µl of Tris-HCl, pH 7.8 was added to neutralize the reaction and the sample was flash frozen until further use. The samples were quantified by liquid scintillation counting (TRI-CARB, PerkinElmer).

## Acknowledgement

We thank NANION Technologies for technical assistance. We thank the staff at the electron microscopy facility at Biozentrum (BioEM lab) and at the PX beamline of the Swiss Light Source (SLS). We thank Tim Sharpe from the Biophysics facility at Biozentrum for SEC-MALS data analysis and Xiaochun Li Blatter for assistance in cell expressions. Work in the Perez lab is supported by the Swiss National Science Foundation (SNSF) (PP00P3_170607), the Helmut Horten Stiftung (HHS), and NANION Research Grant Initiative. Work in the Veening lab is supported by the SNSF (project grants 31003A_172861 and 310030_192517), SNSF JPIAMR grant (40AR40_185533), SNSF NCCR ‘AntiResist’ (51NF40_180541) and ERC consolidator grant 771534-PneumoCaTChER. Work in the Seeger lab is supported by the SNSF (project grant 310030_188817) and a ERC consolidator grant (no. 772190).

## Author Contributions

N.B. performed purifications, crystallization experiments, and functional assays. N.B. and C.P. prepared samples for cryo-EM, processed X-ray and cryo-EM data, built and validated the models. A.R. performed complementation and growth assays under supervision from J.W.-V. G.C. produced samples for sybodies selection. C.A.J.H., and G.C. performed sybodies selection under supervision from M.A.S. G.C. established WTA and LTA extraction assays and N.B. performed extraction experiments. C.P. and N.B. wrote the manuscript with input from all authors. C.P. conceived the project.

## Author Information

### Data and materials availability

Electron microscopy density maps and atomic models have been deposited in the EMDB and PDB, respectively, with accession codes EMD-13268 and PDB 7PAF. Atomic coordinates for the reported crystal structure have been deposited in the PDB under accession code 7B0K.

