## Supplementary_Information for "Mechanistic basis of choline import involved in teichoic acids and lipopolysaccharide modification"

**This PDF file includes:**

Supplementary Text
Figs. S1 to S12
Tables S1

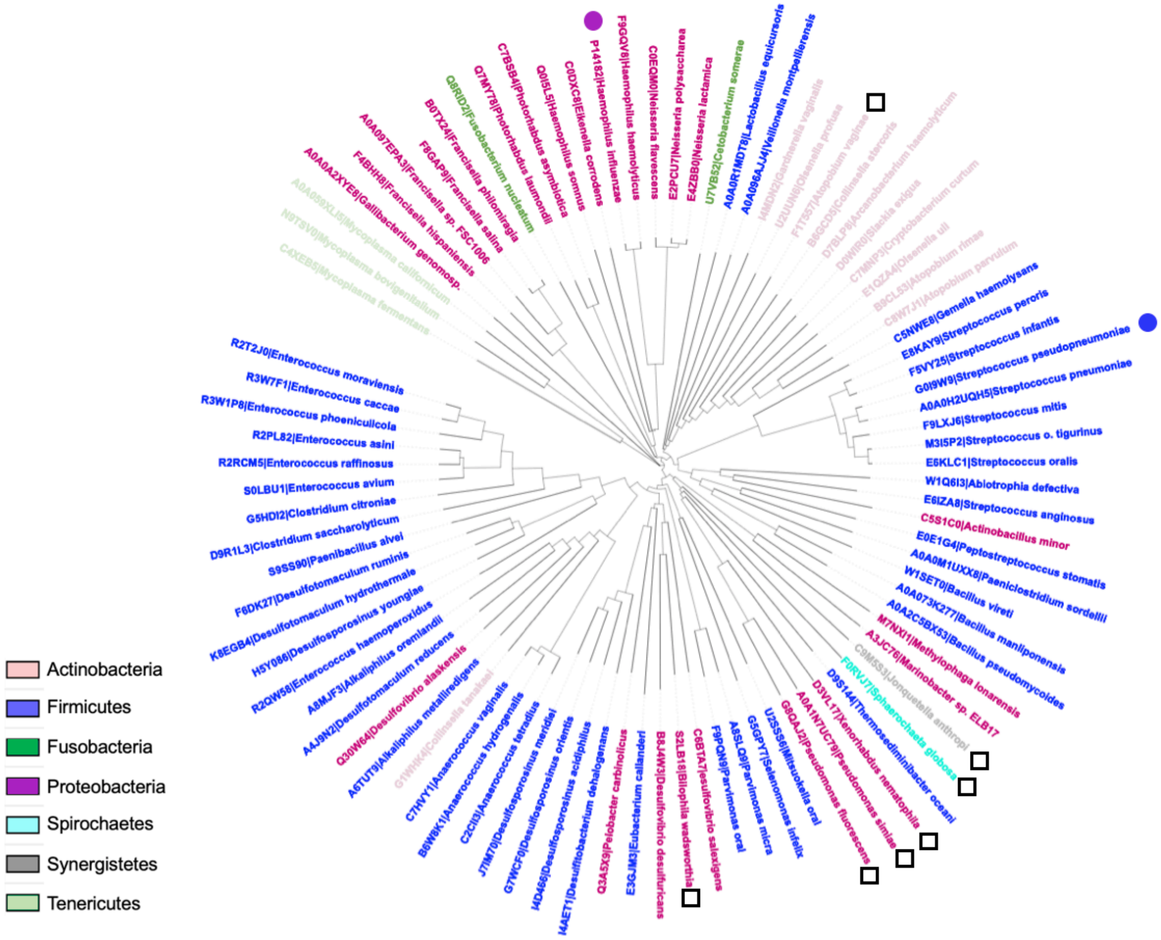

**Fig. S1. LicB is conserved across divergent bacteria phyla.**

Phylogenetic tree of bacteria species across multiple phyla where *lic* operon genes involved in choline uptake (*licB*) and activation (*licA* and *licC*) are conserved. Phyla are depicted by colors according to the inset. *S. pneumoniae* and *H. influenzae* are indicated by blue and magenta dots. White squares show bacteria species where residues E170 and R191 are not conserved (see Fig. 4).

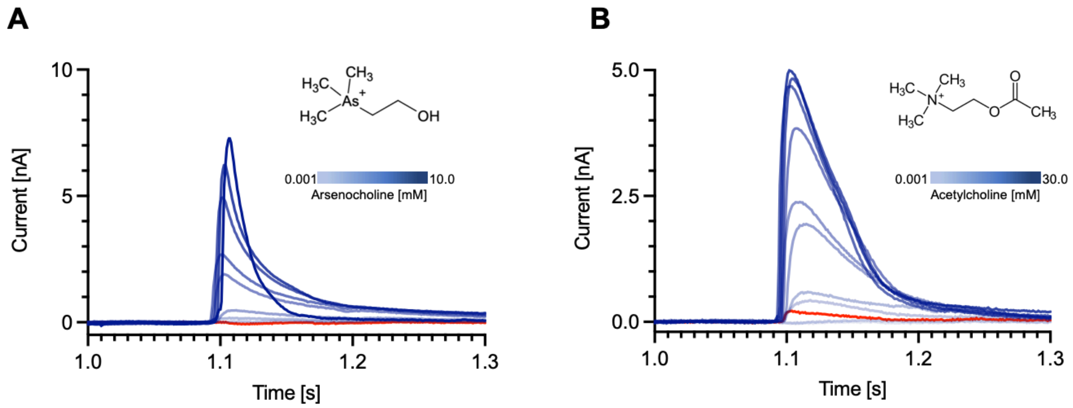

**Fig. S2. LicB displays promiscuous selectivity towards arsenocholine and acetylcholine.**
SSM-electrophysiology recordings of currents measured during application of arsenocholine (**A**) or acetylcholine (**B**) are indicated by blue curves. Protein-free liposomes traces measured in presence of 5 mM arsenocholine (**A**) or 30mM acetylcholine (**B**) are shown in red. The amplitude of the peak currents were used for determination of $EC_{50}$ values as shown in Fig. 2B.

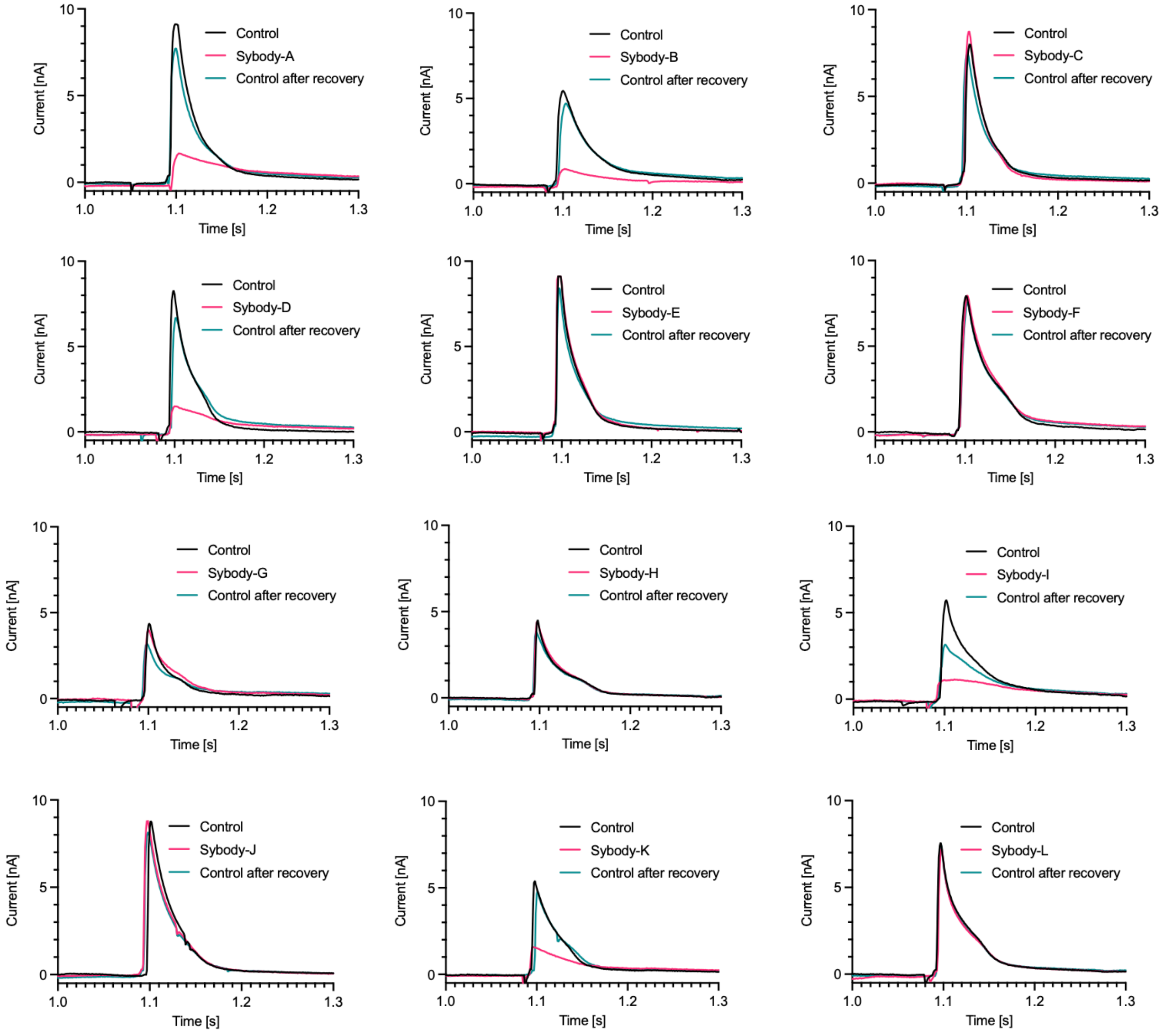

**Fig. S3. SSM-electrophysiology recordings of LicB choline transport in presence of sybodies.** Representative recordings of currents measured during application of 5 mM choline in absence of sybodies are shown in black (control). The same experiment but in presence of 500 nM of sybodies are shown in pink, whereas recordings after unbinding of sybodies are shown in green. The amplitudes of the peak currents (pink traces) were used for the generation of the histogram shown in Fig. 2D.

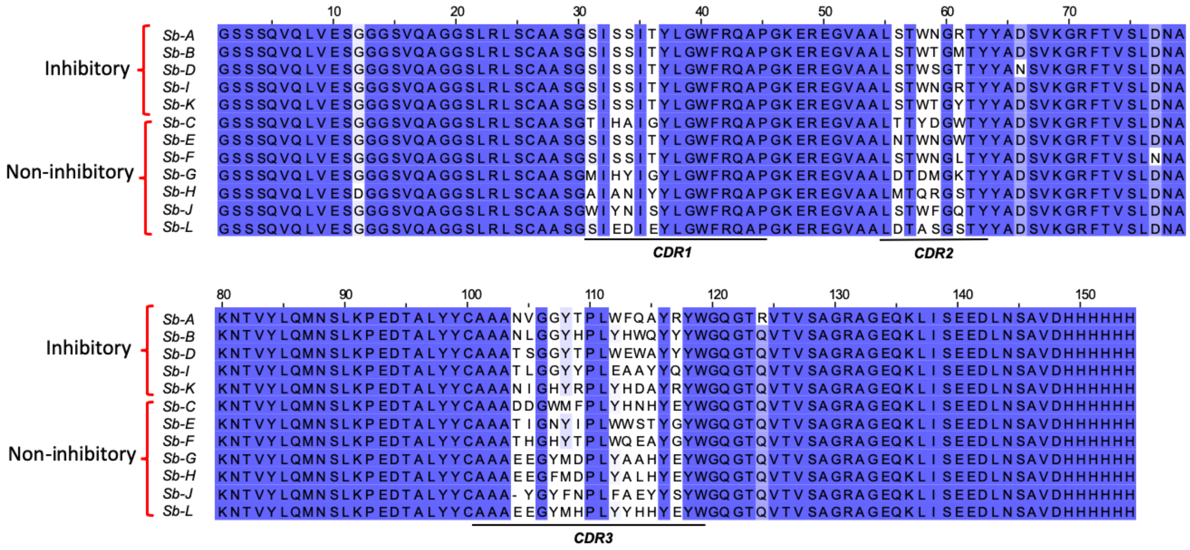

**Fig. S4. Sequence conservation alignment of sybodies characterized by SSM-electrophysiology.**

Inhibitory and non-inhibitory sybodies are grouped together. All sybodies belong to the convex library, which display a long CDR3 region.

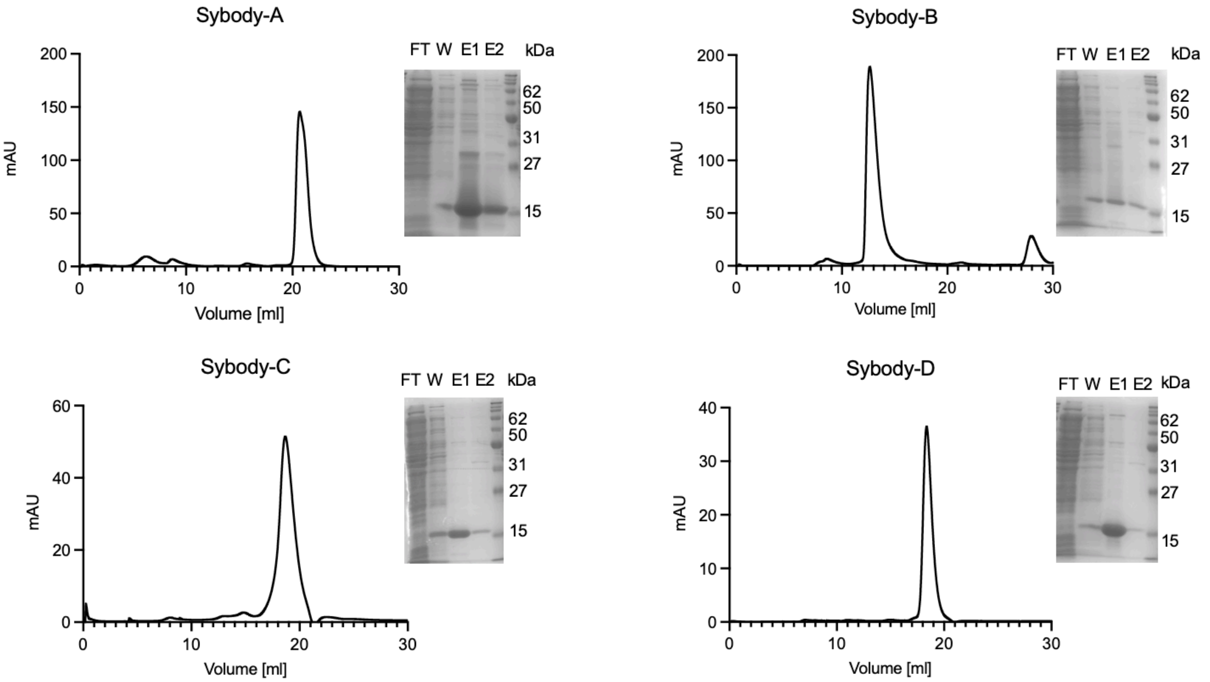

**Fig. S5. Purification of sybodies A, B, C and D.**

(*Left*) Size exclusion profiles using a Superdex 200 Increase 10/300 column. (*Right*) SDS-PAGE of samples from different steps of sybodies purification. FT, flow through. W, washing of column. E1 and E2, two steps elution.

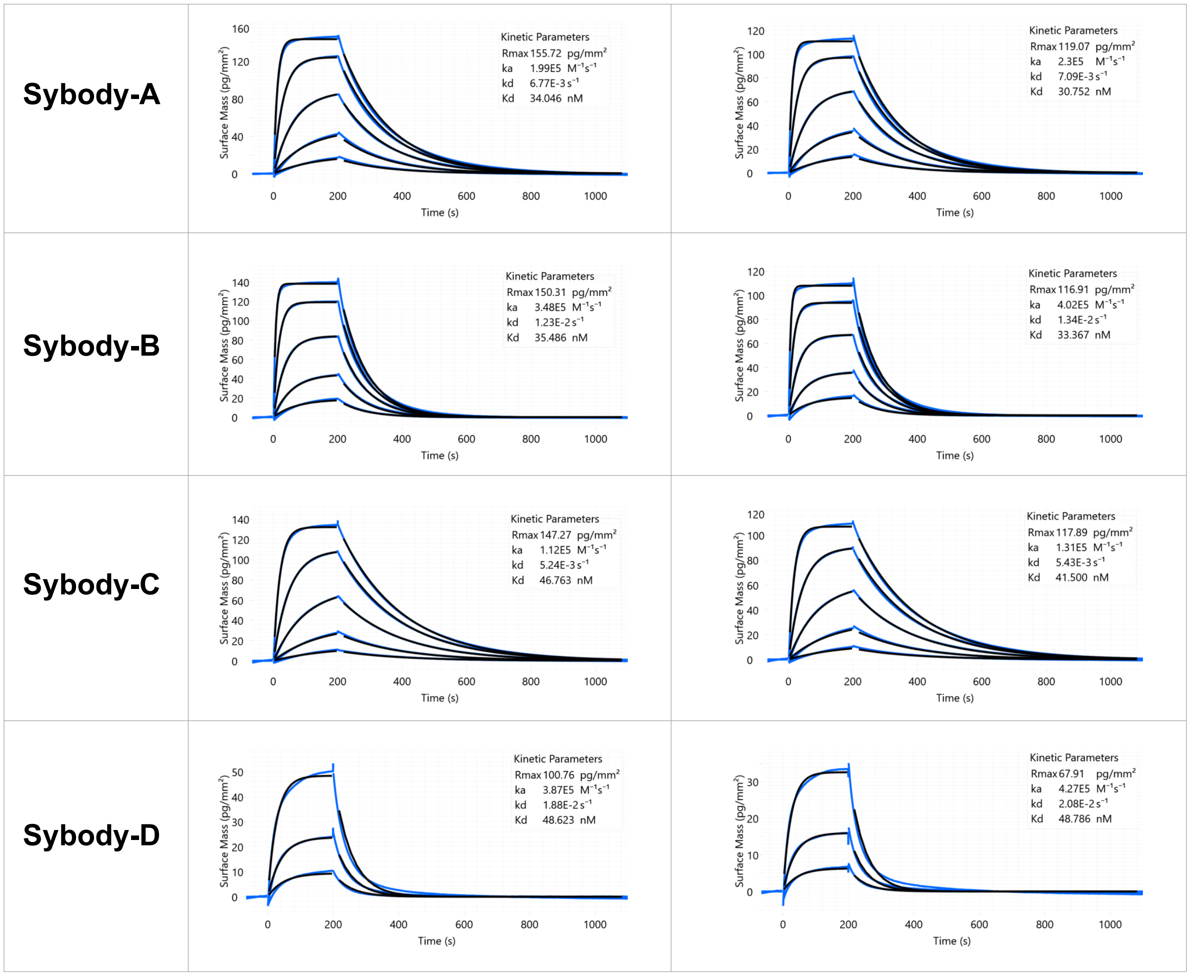

**Fig. S6. Determination of binding affinity of Sybodies to LicB by grating-coupled interferometry (GCI).**

The four sybodies were injected at 5, 15, 45, 135 and 405 nM concentrations in absence (*left*) or presence of 5 mM choline (*right*). Data are shown in blue and fitting curves in black. Data were fitted using a Langmuir 1:1 model.

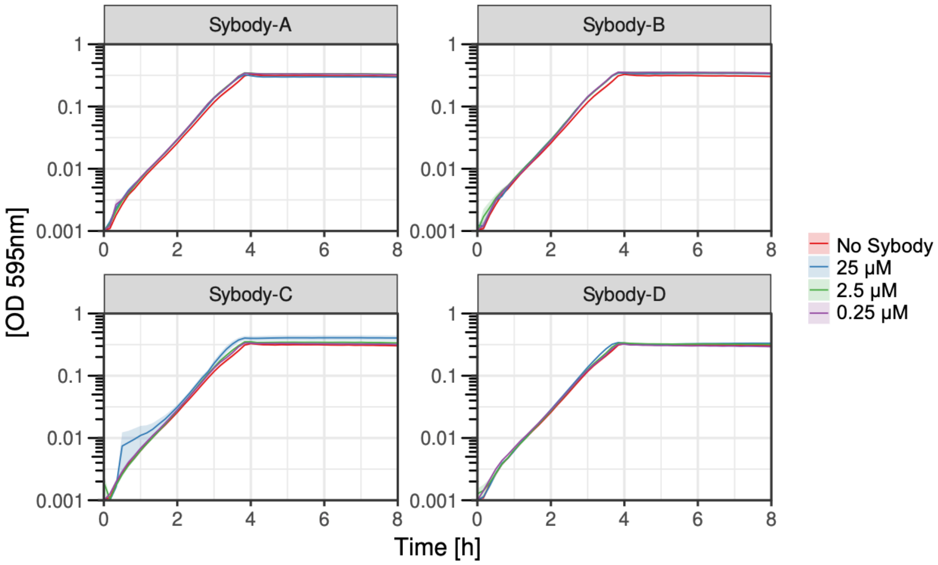

**Fig. S7.** ***S. pneumoniae*** **growth in presence of sybodies.**

Growth curves of unencapsulated *S. pneumoniae* in presence of sybodies targeting LicB in C+Y media supplemented with choline. Three different concentrations of sybodies were tested in comparison with growth without sybody. An average of three replicates and standard error of the mean are plotted.

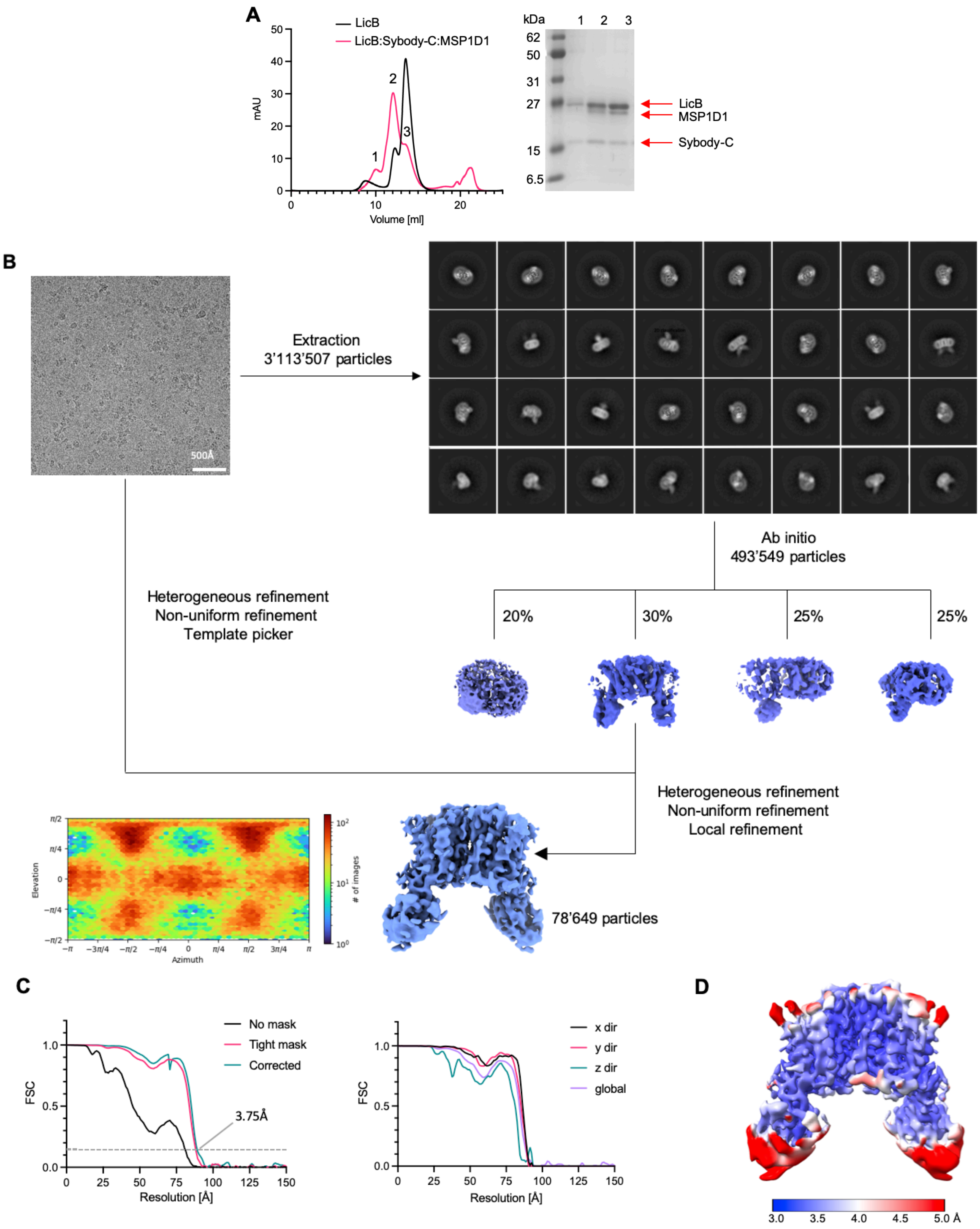

**Fig. S8. Cryo-EM reconstruction of LicB-Sybody-C complex in lipid nanodiscs. A.** Size exclusion chromatography profiles of LicB and LicB:Sybody-C complex reconstituted in nanodiscs in a Superdex 200 Increase 10/300 column. SDS-PAGE of the three main peaks of the red trace is shown. **B**. Representative micrograph of the complex acquired with a Glacios TEM equipped with a K3 camera. The data processing workflow shows 2D classes of LicB:Sybody-C:MSP1D1 after particle extraction and multiple rounds of 2D classification and ab initio reconstruction performed with four classes. The particle classes with best protein-like features were used for further rounds of 3D classification, non-uniform refinement, and template generation. The selected particles were later used for a new round of particle picking. Further processing led to a new optimized set of particles used as an input for iterative heterogeneous, non-uniform, and local refinement rounds with C2-symmetry, which yielded a map at a resolution of 3.75 Å. The heatmap displays the number of particles for a given viewing angle. **C**. (*Left*) FSC plots of masked and unmasked maps calculated by cryoSPARC v.3.2.0. Dashed line indicates a 0.143 cut-off. (*Right*) Directional and global FSC plots calculated by the 3DFSC server. The directional FSC curves providing an estimation of anisotropy of the dataset are shown for directions x, y, and z. **D**. Final 3D reconstruction of the LicB:Sybody-C complex colored according to the local resolution, estimated in cryoSPARC v.3.2.0.

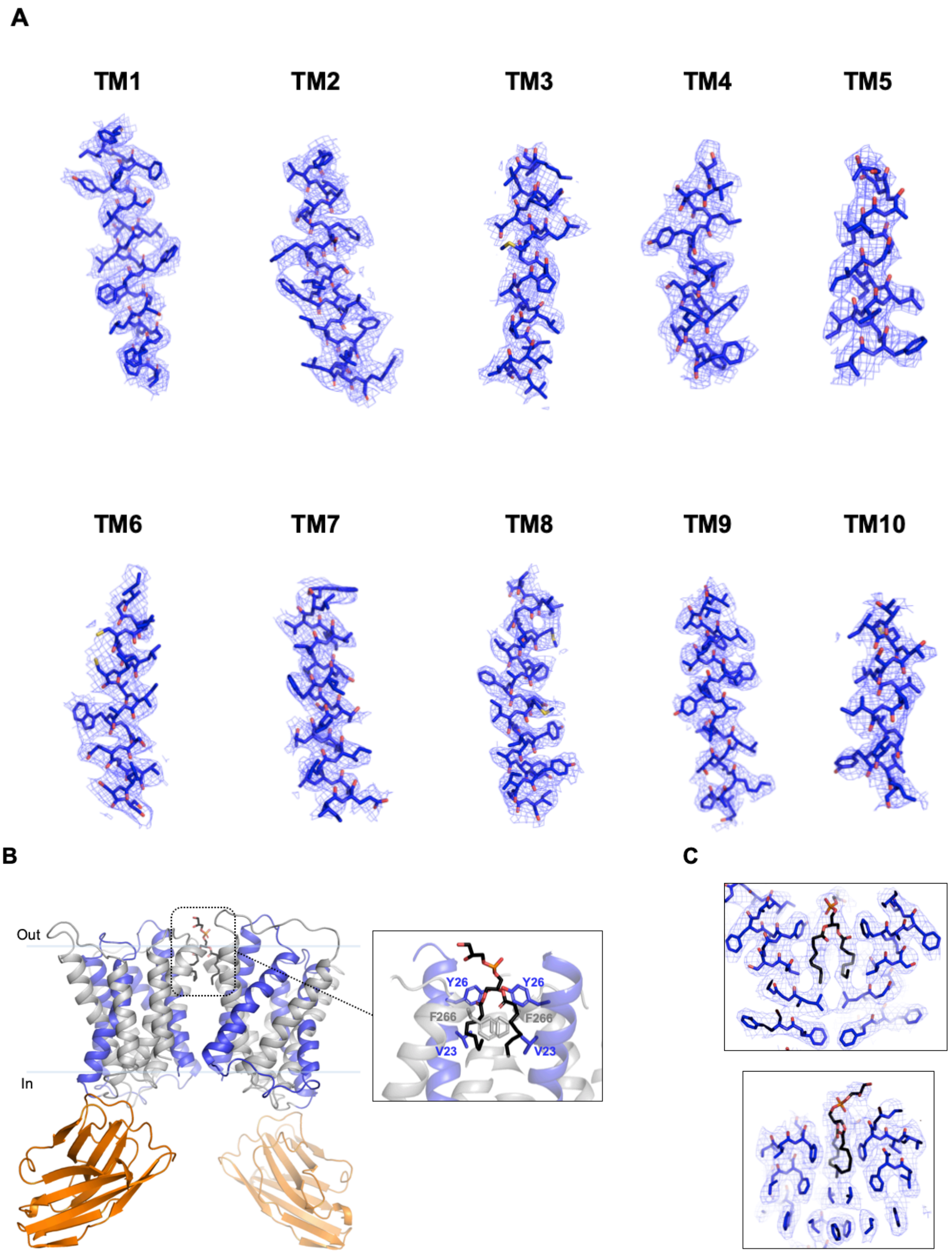

**Fig. S9. Cryo-EM densities of LicB segments.**

**A**. Sections of the cryo-EM density superimposed on the refined structure of LicB. **B**. Side view of the LicB homodimer and bound Sybodies. Inverted repeats TM1-5 and TM6-10 in blue and grey, respectively. Sybody-C is shown in orange. The inset shows a putative POPG lipid molecule present at the dimer interface between both protomers. Surrounding residues are shown. **C**. Cryo-EM density (mesh) superimposed on the POPG lipid and surrounding residues.

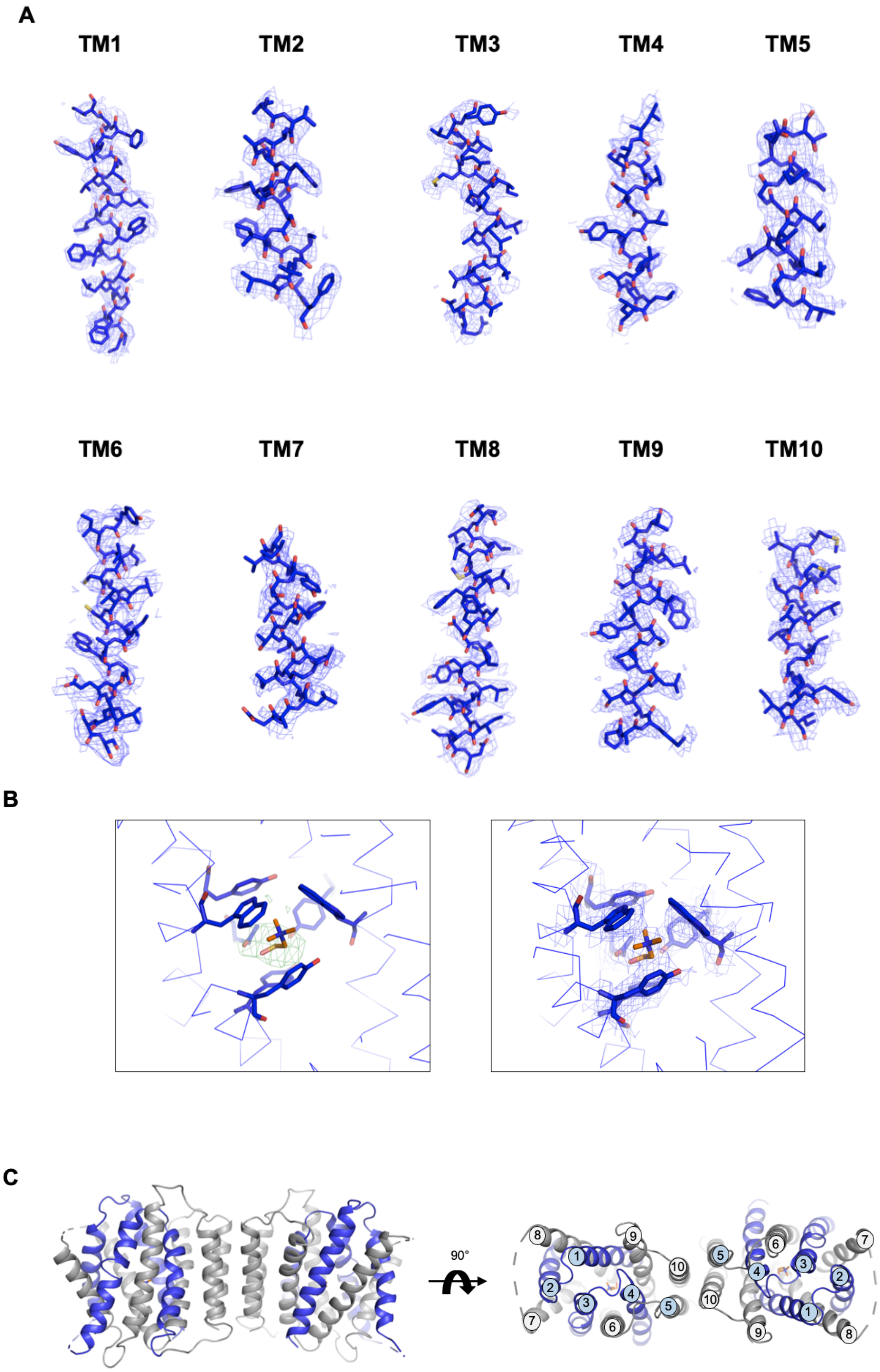

**Fig. S10. Crystal structure of LicB.**

**A**. *2Fo-Fc* electron density map of individual TM segments of LicB at 1.0 σ level. **B**. (*Left*) *Fo-Fc* map at 3.0 σ level showing a positive peak at the choline binding site. (*Right*) *2Fo-Fc* electron density map of choline and coordinating residues after refinement. **C**. Packing of two LicB molecules in the crystal lattice, resembling the arrangement of dimers in other DMT transporters.

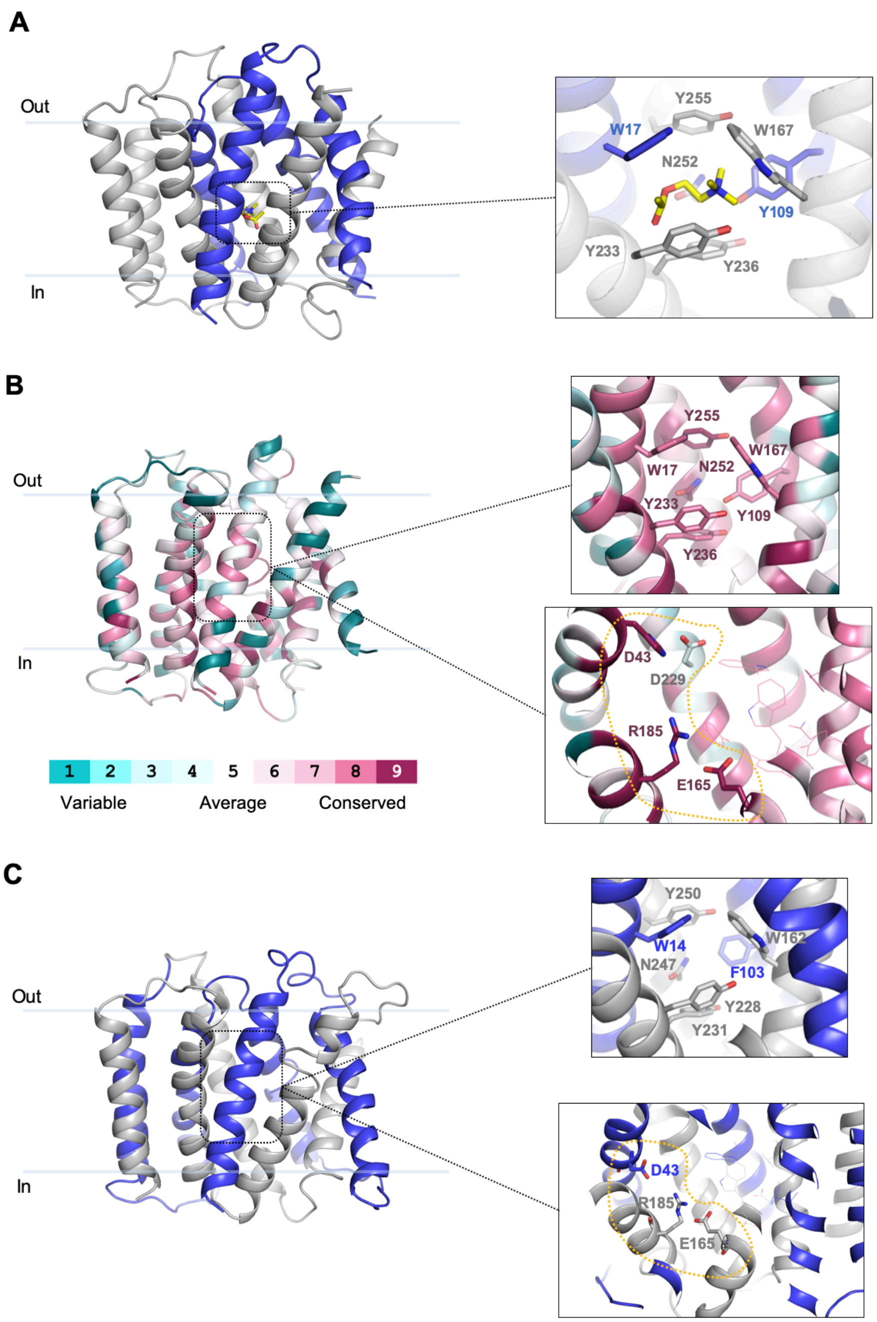

**Fig. S11. Docking of acetylcholine and conservation analysis of residues at the choline binding pocket and surrounding charged residues.**

**A**. Docking of acetylcholine in LicB central cavity. Acetylcholine is shown in yellow. **B**. Sequence conservation analysis. A multiple sequence alignment of LicB homologues sharing more than 35% identity was generated and residues in the LicB structure were colored by sequence conservation (ConSurf server). (*Top*) Residues at the Choline binding site. (*Bottom*) Charged residues close to the choline binding site. C. Homology model of *H. influenza* LicB. (*Top*) Residues at the Choline binding. (*Bottom*) Charged residues close to the choline binding site.

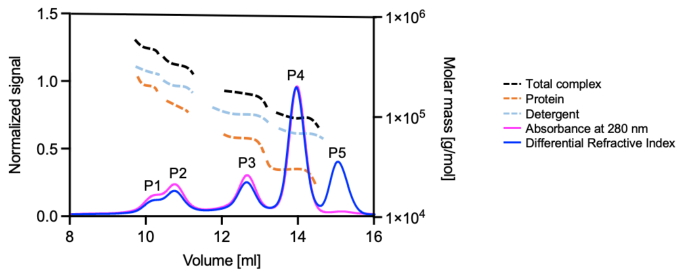

| Peak | Molecular weight [kDa] | | | Interpretation |
|---|---|---|---|---|
| | Total | Detergent | Protein | |
| P1 | 561 | 329 | 232 | Hexamer |
| P2 | 421 | 257 | 164 | Tetramer |
| P3 | 206 | 131 | 75 | Dimer |
| P4 | 125 | 88 | 37 | Monomer |
| P5 | 67 | 66 | 1 | Empty micelles |

**Fig. S12. Size exclusion chromatography with multi-angle light scattering (SEC-MALS) analysis of purified LicB in detergent micelles.**

Size exclusion profile is plotted as normalized signal and MALS apparent molecular masses of the protein in buffer, detergent in buffer and the total complex with dotted lines (right axis). The molecular weights are summarized in the table with the apparent masses of empty micelles (P5), the monomer (P4, main population), dimer (P3), tetramer (P2) and hexamer (P1).

**Table S1.**

Oligos used in this study.

| Strain | Genotype | Reference |
|---|---|---|
| D39V | WT strain of clinical isolate, Serotype 2, parent strain for all strains used in this study unless described | Domenech et al., 2018 |
| VL333 | *prs1::PF6-lacI-tetR (gen)* | Sorg et al., 2020 |
| VL1998 | *prs1::PF6-lacI (gen), bgaA::Plac-dCas9sp (tet)* | Liu et al., 2017 |
| VL4243 | *prsI::PF6-lacI-tetR (gen), bgaA::Plac-licB (tet)* | This study |
| VL4249 | *prsI::PF6-lacI-tetR (gen), bgaA::Plac-licB (tet), zip::Ptet-licB (spc), licB::ery* | Veening collection |
| VL4250 | *prsI::PF6-lacI-tetR (gen), bgaA::Plac-licB-Y233A (tet)* | This study |
| VL4251 | *prsI::PF6-lacI-tetR (gen), bgaA::Plac-licB-Y236A (tet)* | This study |
| VL4252 | *prsI::PF6-lacI-tetR (gen), bgaA::Plac-licB-W17A (tet)* | This study |
| VL4253 | *prsI::PF6-lacI-tetR (gen), bgaA::Plac-licB-W167A (tet)* | This study |
| VL4254 | *prsI::PF6-lacI-tetR (gen), bgaA::Plac-licB-E170A (tet)* | This study |
| VL4255 | *prsI::PF6-lacI-tetR (gen), bgaA::Plac-licB-R191A (tet)* | This study |
| VL4256 | *prsI::PF6-lacI-tetR (gen), bgaA::Plac-licB-H43A (tet)* | This study |
| **Oligo** | **Sequence** | **Reference** |
| R191A_ for | GAAGCCCTCTTAATCGCTCAAGTAACTTCG | This study |
| R191A_ rev | CGAAGTTACTTGAGCGATTAAGAGGGCTTC | This study |
| H43A_ for | GTGGCTGCAACTGCTGATTTTTTGAGCATC | This study |
| H43A_ rev | GATGCTCAAAAAATCAGCAGTTGCAGCCAC | This study |
| Y233A (gBlock) | GATCCTCAATTGCTAGGTCTCATGATTGTTTTTGCAGCCTTTGATATGATT<br>TCCGCGTTGGCTTATTATATCGCTATCAATCGCTTGCAACCAGCCAAGGCT<br>ACAGGCTTGAACGTGAGCTATGTAGTATGGACGGTCTTGTTTGCAGTTGTT<br>TTCTTGGGTGCACCGCTAGATATGCTGACCATTATGACGTCACTTGTCGTC<br>ATTGCTGGAGTTTATATTATTATTAAAGAATAAGGATCCGATCCT | This study |
| Y236A (gBlock) | GATCCTCAATTGCTAGGTCTCATGATTGTTTTTGCAGCCTTTGATATGATT<br>TCCTACTTGGCTGCGTATATCGCTATCAATCGCTTGCAACCAGCCAAGGCT<br>ACAGGCTTGAACGTGAGCTATGTAGTATGGACGGTCTTGTTTGCAGTTGTT<br>TTCTTGGGTGCACCGCTAGATATGCTGACCATTATGACGTCACTTGTCGTC<br>ATTGCTGGAGTTTATATTATTATTAAAGAATAAGGATCCGATCCT | This study |
| N252A (gBlock) | GATCCTCAATTGCTAGGTCTCATGATTGTTTTTGCAGCCTTTGATATGATT<br>TCCTACTTGGCTTATTATATCGCTATCAATCGCTTGCAACCAGCCAAGGCT<br>ACAGGCTTGGCGGTGAGCTATGTAGTATGGACGGTCTTGTTTGCAGTTGTT<br>TTCTTGGGTGCACCGCTAGATATGCTGACCATTATGACGTCACTTGTCGTC<br>ATTGCTGGAGTTTATATTATTATTAAAGAATAAGGATCCGATCCT | This study |
| W17A (gBlock) | GATCCTCATATGAAAAGTAAAAACGGAGTTCCTTTTGGCCTTCTCTCAGGT<br>ATTTTCGCGGGCTTGGGTCTAACGGTTAGTGCTTATATCTTTTCGATTTTTA<br>CAGATTTGTCACCCTTTGTGGTGGCTGCAACTCATGATTTTTTGAGCATCT<br>TTATCTTACTAGCTTTTCTCTTGGTAAAAGAAGGGAAAGTTCGCCTCTCAA<br>TTTTCTTAAATATTCGCAATGTCAGTGTTATCATCGGAGCCTTGCTAGCGA<br>TCCT | This study |
| W167A (gBlock) | GATCCTGCTAGCAGGCCCTATCGGTATGCAGGCCAATCTTTATGCAGTTA<br>AGTATATCGGAAGTTCTTTAGCTTCATCTGTATCGGCTATTTACCCTGCGA<br>TTTCAGTTCTATTGGCTTTCTTCTTTTTGAAGCACAAGATTTCGAAAAATA<br>CTGTATTTGGGATTGTCTTGATTATTGGAGGGATTATTGCTCAGACCTATA<br>AGGTTGAACAGGTTAATTCTTTCTACATTGGGATTCTTTGTGCTTTGGTTT<br>GTGCTATTGCAGCGGGAAGTGAGAGTGTTCTTAGCTCTTTTGCCATGGAA<br>AGTGAATTGAGTGAAATCGAAGCCCTCTTAATCCGTCAAGTAACTTCGTT<br>CTTGTCCTATCTTGTGATTGTGCTCTTCTCTCATCAGTCATTTACTGCAGTA<br>GCCAATGGACAATTGGATCCT | This study |
| E170A (gBlock) | GATCCTGCTAGCAGGCCCTATCGGTATGCAGGCCAATCTTTATGCAGTTA<br>AGTATATCGGAAGTTCTTTAGCTTCATCTGTATCGGCTATTTACCCTGCGA<br>TTTCAGTTCTATTGGCTTTCTTCTTTTTGAAGCACAAGATTTCGAAAAATA<br>CTGTATTTGGGATTGTCTTGATTATTGGAGGGATTATTGCTCAGACCTATA<br>AGGTTGAACAGGTTAATTCTTTCTACATTGGGATTCTTTGTGCTTTGGTTT<br>GTGCTATTGCATGGGGAAGTGCGAGTGTTCTTAGCTCTTTTGCCATGGAA<br>AGTGAATTGAGTGAAATCGAAGCCCTCTTAATCCGTCAAGTAACTTCGTT<br>CTTGTCCTATCTTGTGATTGTGCTCTTCTCTCATCAGTCATTTACTGCAGTA<br>GCCAATGGACAATTGGATCCT | This study |
| OVL2077 | ATTCCTTCTTAACGCCCCAAGTTC | This study |
| OVL2082 | GTCTTCTTTTTTACCTTTAGTAAC | This study |
| OVL5139 | TTAATTCCTTCTTAACGCCCCAAGT | This study |

| | | |
|---|---|---|
| OVL5623 | AGCTGTCGTCTCGATAGATCCTTTCTCCTCTTTAGATCTTTTGAATTCGCG GCCG | This study |
| OVL5624 | AGCTGTCGTCTCGCTATGAAAAGTAAAAACGGAGTTCCTTTTGGCCTTCTC TCAG | This study |
| OVL5625 | AGCTGTCGTCTCGCTTATTCTTTAATAATAATATAAACTCCAGCAATGACG ACA | This study |
| OVL5626 | AGCTGTCGTCTCGtaaGTCGACCTCGAGACTAGTCAAGGTCGGCAATTCTGC AGTA | This study |
| OVL5635 | AACTCTTTCGACAAATGCGCATCGTCTATCTGAAAATAAC | This study |
| OVL5636 | GAAAATGCTATCCAAATGAT | This study |
| OVL6035 | TAAGGGCGTCTCGCGGAAATCATATCAAAGGCTGCAAAA | This study |
| OVL6036 | TAAGGGCGTCTCCTCCGCATTGGCTTATTATATCGCTATCA | This study |
| OVL6037 | AGTGGTCGTCTCGGCAGCCAAGTAGGAAATCATATCAAAGGCTGCAAAA ACAATC | This study |
| OVL6038 | AGTGGTCGTCTCGCTGCATATATCGCTATCAATCGCTTGCAACCAGCCAA GGCTA | This study |
| OVL6039 | GGAATGCGTCTCGGCGAAAATACCTGAGAGAAGGCCAAAAGGAAC | This study |
| OVL6040 | GGAATGCGTCTCGTCGCAGGCTTGGGTCTAACGGTTAGTGCTTAT | This study |
| OVL6041 | GCCATACGTCTCGGCTGCAATAGCACAAACCAAAGCACAGAGAAT | This study |
| OVL6042 | GCCATACGTCTCCCAGCAGGAAGTGAGAGTGTTCTTAGCTCCTTT | This study |
| OVL6043 | CATCCACGTCTCGGCACTTCCCCATGCAATAGCACAAACCAAAGC | This study |
| OVL6044 | CATCCACGTCTCCGTGCAAGTGTTCTTAGCTCCTTTGCTATGGAA | This study |
| OVL6045 | GACAACCGTCTCCGCGATTAAGAGGGCTTCGATTTCACTCAGTTC | This study |
| OVL6046 | GACAACCGTCTCCTCGCACAAGTGACTTCGTTCTTGTCCTATCTT | This study |
| OVL6047 | CTCCTACGTCTCCGCAGTTGCAGCCACCACAAAGGGTGACAAATC | This study |
| OVL6048 | CTCCTACGTCTCCCTGCAGATTTTTTGAGCATCTTTATCTTACTA | This study |
